# Expression levels of the Band-7 protein FLOTILLIN modulate salt tolerance, growth and development in the moss *Physcomitrium patens*

**DOI:** 10.1101/2025.04.14.648360

**Authors:** Erika Csicsely, Norina Noor, Susanne Mühlbauer, Hans-Henning Kunz, Serena Schwenkert, Martin Lehmann, Andreas Klingl, Oguz Top, Wolfgang Frank

## Abstract

The Band-7 proteins, known as FLOTILLINs (FLOT), are present at the plasma membranes of most land plants. They function in clathrin-independent endocytosis and contribute to nodule formation following symbiotic infections. This study reveals that the single FLOT variant in *Physcomitrium patens* is located at the thylakoid membranes in chloroplasts, serving an unanticipated function. Phenotypic analysis of knockout and overexpression lines demonstrates that *PpFLOT* overexpression significantly impairs the high salinity tolerance of *P. patens*. Additionally, liquid protonema cultures of *PpFLOT-*OEX lines exhibited a distinct color change due to necrotic events and developed brachycyte-like cells. These changes correlate with the strength of *PpFLOT* expression and do not occur when these lines are cultivated on solid medium. Our study found that *PpFLOT*-OEX lines display increased chlorophyll and H_2_O_2_ production. We also discovered that PpFLOT is regulated by ABA and light, and its high expression can potentially affect retrograde signaling. Metabolomics and proteomics analyses revealed changes in the pigment and lipid composition as well as differentially accumulated proteins in *PpFLOT* mutant lines. We also observed changes in the expression of ion-transport related genes, accumulation of lipids crucial during pathogen defense, and differentially accumulated proteins taking part in multiple metabolomic pathways. Consequently, our study suggests a novel role for chloroplastic PpFLOT in plant terrestrialization, as it is putatively involved in Ca^2+^ and reactive oxygen species (ROS) signaling in response to abiotic and biotic stress, along with the light-dependent regulation of chlorophyll biosynthesis.

## Introduction

The Stomatin/Prohibitin/Flotillin/HflK/C (SPFH) protein family, also known as Band-7 proteins, is widespread across evolutionary lineages (Rivera-Milla et al., 2006; Daněk et al., 2016; Martiniere and Zelazny, 2021). The SPFH-protein family includes FLOTILLINS (FLOT), STOMATINS (STOM), PROHIBITINS (PHB), ERLINS (ER), the bacterial membrane-specific HflK and HflC proteins, and the plant-specific HYPERSENSITIVE-INDUCED REACTION proteins (HIR). Despite differences in function, these proteins share the SPFH/Band-7 domain, ensuring cell membrane-associated localization, and a common accumulation in micro/nanodomains in cell membranes (Daněk et al., 2016; Martiniere and Zelazny, 2021). Enriched in detergent-resistant membrane fractions (DRM), these SPFH-proteins are proposed to actively participate in microdomain formation (Browman et al., 2007; Martiniere and Zelazny, 2021). Among these SPFH-proteins, FLOT seems to have evolved independently in multiple lineages since plant and fungi FLOT do not encode a Flotillin domain or a PDZ3-binding motif that are characteristic for metazoan flotillin/reggie proteins (Rivera-Milla et al., 2006). A recent study suggests a horizontal gene transfer from fungi to plants, indicating FLOTś role in the infection of nitrogen-fixing bacteria, endocytosis, and seedling development evolved after this transfer (Ma et al., 2022). In plants, FLOT is sterol-dependently recruited into nanodomains, as demonstrated by altered dynamics in response to sterol-depleting agents like methyl-β-cyclodextrin (mβCD) and disturbances in sterol biosynthesis like in *cyclopropylsterol isomerase 1* (*cip-1*) mutant lines of *Arabidopsis thaliana* which resulted in modified FLOT1 distribution in the root tissue (Li et al., 2012; Cao et al., 2020; Martiniere and Zelazny, 2021). Studies on FLOT1 in *A. thaliana* revealed its crucial role in clathrin-independent endocytosis (Li et al., 2012; Hao et al., 2014; Daněk et al., 2016). AtFLOT2 interaction partners in *A. thaliana*, including aquaporins and carbonic anhydrase 2, an early-responsive dehydration stress protein, have essential functions in biotic and abiotic stress response (Junková et al., 2018). Additionally, both FLOT2 and FLOT4 of *Medicago truncatula* are critical in the early stages of symbiotic bacterial infection, with the loss of these genes impairing nodule formation induced by *Sinorhizobium meliloti* infection (Haney and Long, 2010). Furthermore, Kroumanová et al. (2019) demonstrated increased expression of all three *AtFLOT* variants in response to Flagellin 22 (flg22) peptide treatment, while salt treatment suppressed *AtFLOT1* and *AtFLOT2* expression in *A. thaliana*. Notably, flg22 treatment increased trafficking of AtFLOT1 into late endosomes, suggesting its advantageous degradation during pathogen response (Yu et al., 2017).

Suppression of *AtFLOT1* expression in *A. thaliana* leads to reduced seedling growth and root length (Li et al., 2012), while silencing *MtFLOT2*, *MtFLOT3*, and *MtFLOT4* in *M. truncatula* alters root development (Haney and Long, 2010). Compared to *A. thaliana*, which encodes three FLOT variants, AtFLOT1 (AT5G25250), AtFLOT2 (AT5G25260), and AtFLOT3 (AT5G64870), *Physcomitrium patens* encodes only one (PpFLOT; Pp3c3_21910). This PpFLOT shares significant protein sequence similarity (73 – 75 %) with each of the three AtFLOT variants (Phytozome v.13; https://phytozome-next.jgi.doe.gov). Since *P. patens* is a model organism for the land plant adaptation, the unique expression of a single *FLOT* compared to the multiple variants in seed plants, suggests an evolutionary conserved role potentially linked to plant terrestrialization. Notably, our recent study disrupting a *DICER-LIKE1a* (*PpDCL1a*) autoregulatory feedback loop based on intronic microRNA (miRNA) processing not only increased *PpFLOT* expression levels, but also led to salt sensitivity and an ABA hyposensitivity (Arif et al., 2022). This indicates miRNA-mediated expression control of *PpFLOT* and its role in the abiotic stress tolerance in *P. patens.* Our analysis of Δ*PpFLOT* and *PpFLOT*-OEX lines reveals PpFLOT as a negative regulator of salt stress response, contrasting with its potentially positive role in infection events and the biotic stress response based on our proteomics analysis. In-depth investigations via ‘omics’ approaches revealed alterations in the accumulation of proteins, lipids, and pigments, impacting salt stress tolerance and further suggesting a potential role in pathogen infection. Localization studies indicated PpFLOT association with thylakoid membranes, which contrasts its subcellular localization prediction as well as the localization of FLOTs in other plant species that are localized to the plasma membrane. Changes in protein and lipid profiles further support the involvement of PpFLOT in multiple chloroplast-related metabolic pathways, linking it to both salt stress response and pathogen defense.

## Results

### Localization of single FLOT variant in *P. patens*

Even though all SPFH-proteins are associated with cell membranes, their association is notably specific and determined by the N-terminal region of the protein. The N-terminus of SPFH-proteins, housing transmembrane domains or hydrophobic regions in conjunction with the Band-7 protein domain, facilitates interaction with cell membranes (Rivera-Milla et al., 2006; Browman et al., 2007; Daněk et al., 2016). Previous studies in human cells have demonstrated that the membrane association of FLOT is determined by its N-terminal region (Daněk et al., 2016). Localization studies in *A. thaliana* identified all three FLOT variants in the plasma membrane of root epidermal cells (Li et al., 2012; Danek et al., 2020), and for AtFLOT1 and AtFLOT2 in epidermal cotyledon cells (Junková et al., 2018; Cao et al., 2020). AtFLOT1 was also detected in the tonoplast of root epidermal cells (Danek et al., 2020), most likely due to its primary role in clathrin-independent endocytosis (Li et al., 2012). To determine the localization of FLOT in *P. patens* we constructed a *PpFLOT* CDS construct with an added citrine tag in-frame by cloning it into an empty Actin 5 (ACT5) vector containing the linked citrine sequence (Top et al., 2021). *P. patens* WT protoplasts were transformed with this construct to generate transient *PpFLOT::citrine* lines. Three days after transformation the localization of PpFLOT::citrine was traced by confocal microscopy. In contrast to all three FLOT variants in *A. thaliana,* PpFLOT::citrine did not localize to the plasma membrane of the protoplast, but clearly colocalized with chlorophyll autofluorescence signals in the chloroplasts (Figure 1A). Close-up images further disclosed the accumulation of PpFLOT::citrine in thylakoids, suggesting the formation of nanodomains similar to the accumulation of AtFLOTs in plasma membranes. *In silico* prediction tools did not indicate chloroplast localization for PpFLOT, underscoring the limitations of computational approaches in predicting subcellular protein distribution. To determine whether PpFLOT exhibits a similar localization pattern in other plant species, we cloned the *PpFLOT* CDS into the pHKL0786 vector containing a venus tag and transiently expressed it in *Nicotiana benthamiana* leaves. Confocal microscopy analysis three days post-transformation revealed that PpFLOT::venus predominantly localized to the plasma membrane, consistent with FLOT localization in *A. thaliana* (Li et al., 2012; Daněk et al., 2016). Notably, in contrast to *P. patens*, no detectable fluorescence signal was observed in the chloroplasts, indicating that PpFLOT does not associate with plastids in tobacco cells (Supplementary Figure 1).

**Figure 1:**
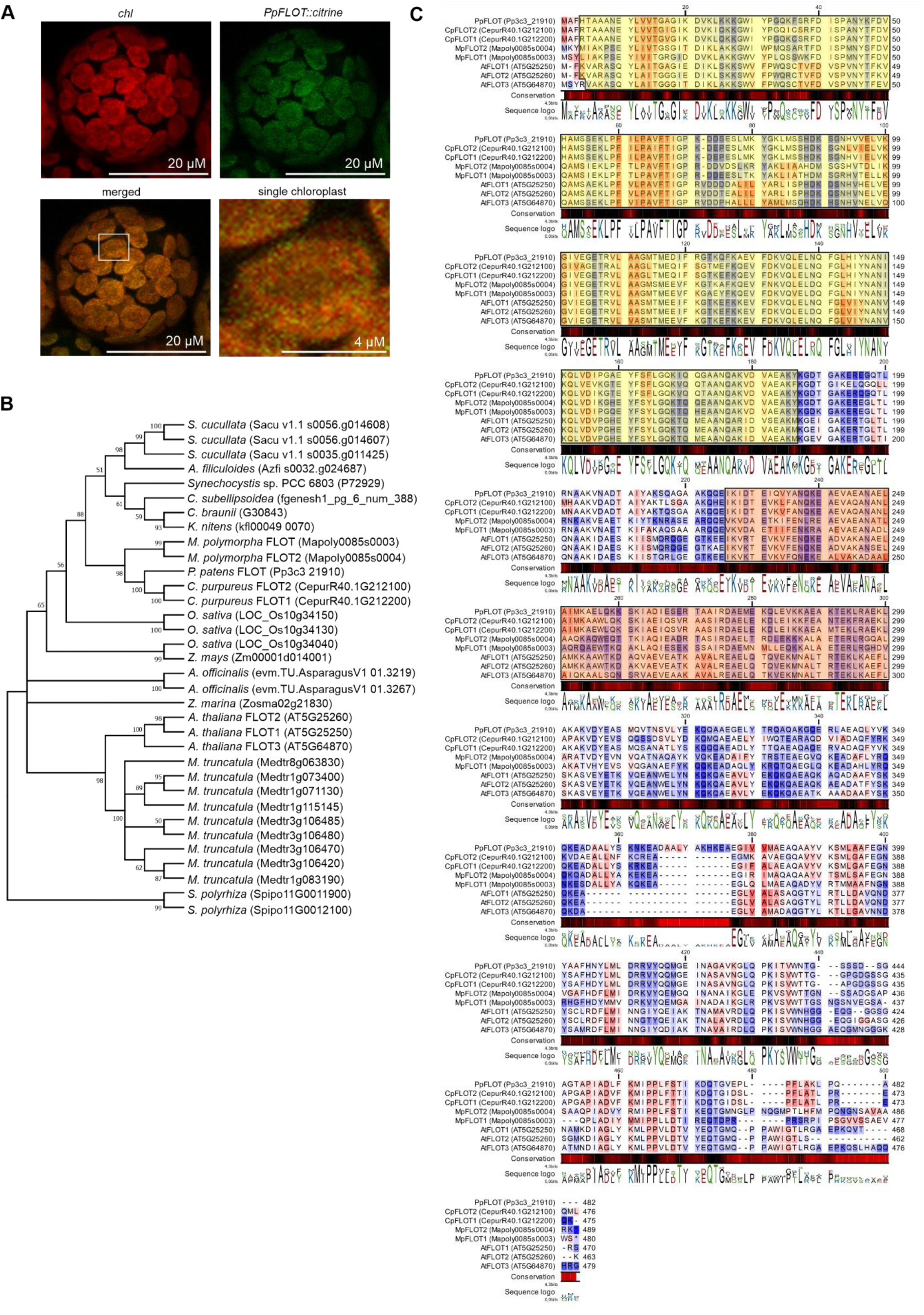
Localization of PpFLOT::citrine and comparison of its peptide sequence to homologs in other plant species. (A) Confocal microscopy images of a *P. patens* protoplast transiently transformed with the *PpFLOT::citrine* construct showing chlorophyll (chl) (red), *PpFLOT::citrine* (green), and a merged (orange) image. A close-up of a single chloroplast of the merged image is also presented. Scale bars are indicated in the respective images. (B) Phylogenetic tree generated by MEGA X using the maximum likelihood method, depicting relationships among PpFLOT homologs in various plant species: *A. officinalis*, *A. thaliana*, *A. filiculoides*, *C. purpureus*, *C. braunii*, *C. subellipsoidea*, *K. nitens*, *M. polymorpha*, *M. truncatula*, *O. sativa*, *P. patens*, *S. cucullata*, *S. polyrhiza*, *Synechocystis* sp. PCC 6803, *Z. mays* and *Z. marina*. The alignment was generated with the CLC workbench v20.0.4. Only branches with bootstrap values > 50 are shown. (C) Protein sequence alignment of all FLOT variants from *A. thaliana*, *C. purpureus*, *P. patens,* and *M. polymorpha* generated by CLC workbench v20.0.4. Sequence conservation is color-coded, from 0 % (red) to 100 % (black), and a sequence logo is provided below the alignment. Protein sequence hydropathicity is marked according to Kyte-Doolittle (Kyte and Doolittle, 1982) from minimal (blue) to maximal (red). SPFH/Band-7 protein domain regions are highlighted in yellow and regions of potential coiled-coil structures are highlighted in orange.

### Analysis of the evolutionary relationship and the domain structure of PpFLOT

Phylogenetic analysis of PpFLOT was conducted through reciprocal BLAST of full-length peptide sequences from various plant species, *Asparagus officinalis*, *A. thaliana*, *Azolla filiculoides*, *Ceratodon purpureus*, *Chara braunii*, *Coccomyxa subellipsoidea*, *Klebsormidium nitens*, *Marchantia polymorpha*, *M. truncatula*, *Oryza sativa*, *P. patens*, *Salvinia cucullata*, *Spirodela polyrhiza*, *Synechocystis* sp. PCC 6803, *Zea mays*, and *Zostera marina*. Results indicated a high similarity between bryophyte FLOT homologs, with PpFLOT showing a closer relationship to homologs in ferns, green algae and cyanobacteria than to seed plant homologs (Figure 1B). Notably, PpFLOT homologs in monocotyledons (*O. sativa*, *Z. mays*, *Z. marina* and *A. officinalis*) exhibit higher similarity compared to homologs in dicotyledonous species like *A. thaliana* and *M. truncatula* (Figure 1B). Interestingly, both *O. sativa* and *Z. marina*, which grow submerged or partially submerged in water, share higher similarity with PpFLOT. Given *P. patens‵* ability to be cultivated in submerged conditions, this suggests that PpFLOT’s function may have evolved during the water-to-land transition and remains beneficial for plants regularly encountering anoxic environments.

To examine the potential impact of amino acid (aa) sequence variations on the distinct locations of PpFLOT and its *A. thaliana* variants full-length homologs of PpFLOT were identified through reciprocal BLAST searches in *A. thaliana*, *C. purpureus*, and *M. polymorpha*. Alignment and analysis using CLC workbench v20.0.4 (Qiagen) revealed minimal changes in the aa sequence, with a generally high conservation among the FLOT variants, except for the last 100 aa at the C-terminus. Additionally, PpFLOT exhibited 22 aa from a repeated DAALY*K*KEA motif at positions 354 to 375 (Figure 1C). This conserved motif, also present in related bryophyte species, was absent in all three AtFLOT variants (Figure 1C). These minimal changes within an otherwise conserved sequence support a common origin of FLOT across all plant species. Evolutionary alterations in the FLOT peptide sequence might have occurred in the last common ancestor of bryophytes and tracheophytes, potentially resulting in an altered function of FLOT in bryophytes. The analyzed proteins encode an SPFH/Band-7 protein domain in the N-terminal region, beginning at position three, or position four in AtFLOT3, and ending at position 185/6 (Figure 1C). This region, crucial for membrane association, contains hydrophobic regions, displaying slight changes in hydrophobicity between bryophyte FLOT and the three *A. thaliana* FLOTs (Figure 1C). Another characteristic feature of FLOTs are coiled-coil domains that serve as recognition sites for interaction partners and facilitate oligomerization with themselves or other FLOT variants at the plasma membrane (Rivera-Milla et al., 2006; Frick et al., 2007; Solis et al., 2007; Daněk et al., 2016). Interaction studies with truncated AtFLOT1/3 proteins in yeast demonstrated oligomerization between AtFLOT1 and AtFLOT3 (Yu et al., 2017). These studies showed that AtFLOT1 aggregated at plasma membranes (Yu et al., 2017). Further confirmation of the crucial role of coiled-coil regions in oligomerization was provided when truncated AtFLOT1/3 variants lacking these regions (position 201 - 470) (Figure 1C) failed to interact (Yu et al., 2017). Using the Galaxy-based (ver. 5.0.0.1) (Galaxy community, 2022) application pepcoil (Rice et al., 2000; Blankenberg et al., 2007) and the web-based application CoCoNat (Madeo et al., 2023), a search for coiled-coil regions in PpFLOT identified putative structures between positions 226 to 300 (Figure 1C). Given the presence of these structures, it is likely that PpFLOT undergoes homo-oligomerization, leading to the formation of PpFLOT scaffolds in thylakoid membranes, providing anchoring points for protein complex formation and interactions.

### Generation of Δ*PpFLOT* knockout and overexpression lines

To probe the potential role of PpFLOT in *P. patens* development, we generated Δ*PpFLOT* lines by replacing exon 2 of the *PpFLOT* CDS with a *nptII* selection cassette via homologous recombination (Figure 2A) (Schaefer, 2001; Frank et al., 2007). Following protoplast transformation with the linearized construct and subsequent selection via G418 sulfate (50 µg/ml), transformed plants were screened by amplifying the complete genomic *PpFLOT* sequence (Figure 2B). Examination of 3’ site and 5’ site integration of the construct (Figure 2B), along with confirming *PpFLOT* transcript loss (Figure 2C), identified three independent Δ*PpFLOT* lines, Δ*PpFLOT-1,* Δ*PpFLOT-2* and Δ*PpFLOT-3*, with a single integration confirmed by Southern blot analysis (Figure 2D). Given our previous observation of increased *PpFLOT* expression linked to salt sensitivity and ABA hyposensitivity in *P. patens* (Arif et al., 2022), we also generated *PpFLOT* overexpression lines. To generate the *PpFLOT-*OEX lines we prepared a construct in which the *PpFLOT* CDS was under the control of an *Act5* promoter for constitutive expression and the respective also harbored a hygromycin resistance cassette for plant selection. After protoplast transformation, screening and confirmation of construct insertion in the genome, an enhanced expression was confirmed by amplifying the *PpFLOT* transcript from cDNA for 25 and 35 cycles by PCR (Figure 2E). This approach identified three *PpFLOT-*OEX lines, *PpFLOT*-OEX1, *PpFLOT*-OEX2 and *PpFLOT*-OEX3 that showed a relative expression of 202.79 (SEM ± 62.01), 317.61 (SEM ± 102.18) and 674.49 (SEM ± 98.6), respectively, compared to the WT and normalized against the expression of *ELONGATION FACTOR 1 ALPHA* (*PpEF1α*) (Figure 2F).

**Figure 2:**
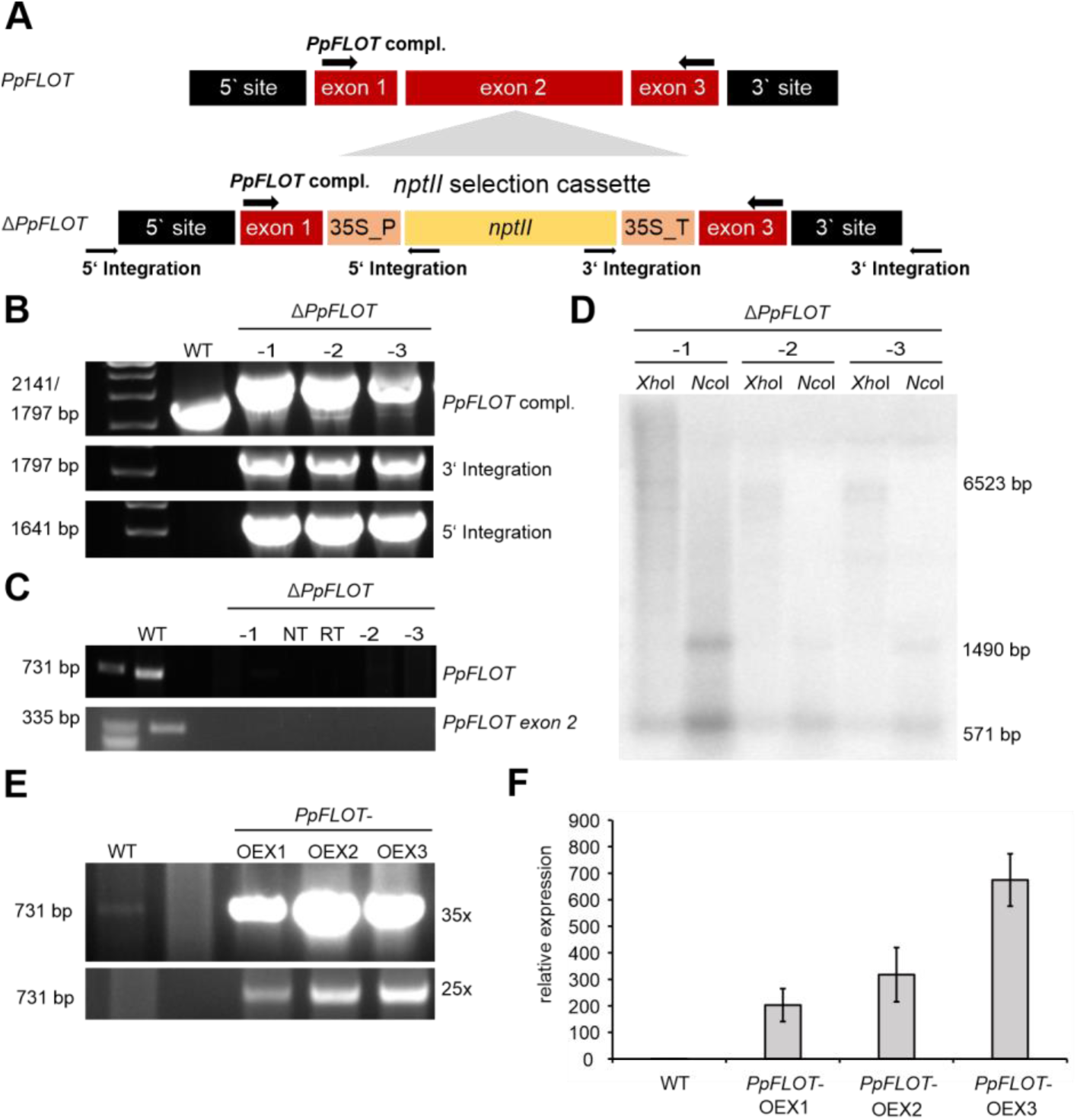
Generation of Δ*PpFLOT* and *PpFLOT*-OEX lines. (A) Schematic representation of the transformation construct. The upper panel illustrates the WT *PpFLOT* coding sequence (CDS), while the lower panel displays the construct for Δ*PpFLOT* generation, involving exon 2 replacement with a *nptII* selection cassette. Arrows indicate primer positions during screening (B) Screening of Δ*PpFLOT* lines via PCR, amplifying the complete *PpFLOT* sequence and the 5’ and 3’ integration sites using genomic DNA (C) Transcript analysis of *PpFLOT* by PCR from cDNA, amplifying both the *PpFLOT* transcript sequence and exon 2 sequence. (D) Southern blot analysis for all three identified Δ*PpFLOT* lines, confirming single construct integration in the moss genome. In case of single integration, digestion of total genomic DNA with *Xho*I and detection of the *nptII* selection cassette with a complementary probe results in a band of 6523 bp since *Xho*I does not cut within the selection marker sequence. For better validation, the experiment was repeated with *Nco*I, an enzyme that cuts within the *nptII* selection cassette and produces two bands (1490 bp and 571 bp) upon single integration. (E) Confirmation of *PpFLOT*-OEX lines by PCR amplification of *PpFLOT* transcript from cDNA at 35 and 25 cycles, comparing band intensities to WT (F) qRT-PCR of relative *PpFLOT* expression compared to the WT, normalized to *PpEF1α* expression, following Schmittgen and Livak (2008). Mean values with error bars indicating ± SEM (n = 3) are presented. Oligonucleotide sequences are listed in Supplementary Table 1.

### Δ*PpFLOT* lines display an enhanced salt tolerance

Liquid cultures of *P. patens* WT and all three generated Δ*PpFLOT* lines were grown for 14 d under control conditions and in medium supplemented with 250 mM NaCl. To detect potential changes in the accumulation of biomass and their salt sensitivity every two to three days, the dry weight of the equalized cultures was determined, and images of the cultures were taken.

Despite reported stunted growth in AtFLOT1 amiRNA lines (Li et al., 2012), no changes in accumulated biomass over time were observed in the Δ*PpFLOT* lines compared to the WT, both under control conditions and in salt-treated cultures (Figure 3A, 3B). Interestingly, unlike the WT, the Δ*PpFLOT* lines did not display bleaching in response to the salt treatment. Bleaching in plant cultures typically results from a decrease in chlorophyll content (Marconi et al., 2001; Taïbi et al., 2016) to avoid reactive oxygen species (ROS) accumulation and oxidative damage (Verma and Mishra, 2005). The apparent lack of chlorophyll suppression in response to salt treatment suggests a potential role for PpFLOT in detecting or regulating ROS, or in chlorophyll biogenesis regulation, a function that may be impaired in Δ*PpFLOT* lines.

**Figure 3:**
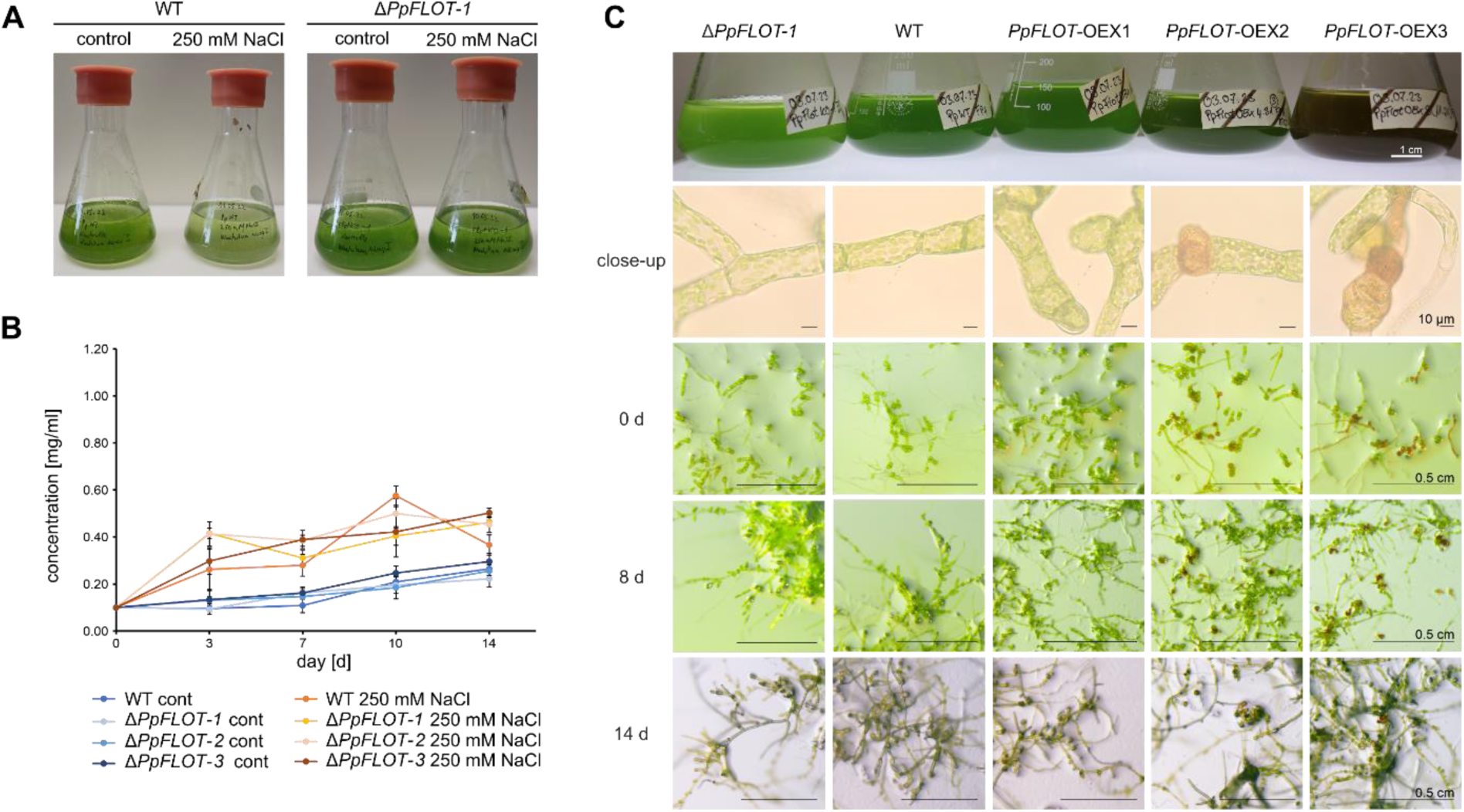
Growth phenotype of Δ*PpFLOT* and *PpFLOT*-OEX lines. (A) Comparison of WT and Δ*PpFLOT-1* liquid protonema cultures with an initial density of 0.1 mg/ml after 14 d of treatment with 250 mM NaCl (right) and an untreated control (left). (B) Growth curves of Δ*PpFLOT* lines and WT determined by dry weight measurement every three to four days for 14 d. Cultures were initially inoculated with an equal density of 0.1 mg/ml dry weight. Measurements were taken during treatment with 250 mM NaCl and under control conditions for Δ*PpFLOT* lines and a WT control. (C) Upper panel displays liquid protonema cultures starting with an equal density of 100 mg/L and grown for 16 weeks (left to right) of Δ*PpFLOT-1*, WT, *PpFLOT*-OEX1*, PpFLOT*-OEX2, and *PpFLOT*-OEX3 at a density of ∼ 1 mg/ml dry weight. The second panel presents close-ups of protonema cells from the respective lines when cultivated in standard liquid medium. Diaspore-like cells with brown coloration develop in *PpFLOT*-OEX2 and 3. Panels three to five show the development of the respective lines when cultivated for 0, 8 and 14 d on solid growth medium. Scale bars are indicated in the respective images.

### Increased *PpFLOT* overexpression strongly affects protonema growth

Long-term cultivation of liquid protonema cultures from all generated *PpFLOT* mutant lines (including Δ*PpFLOT-1* and *PpFLOT*-OEX1,2,3 lines) resulted in changes of the coloration. In Figure 3C depicted cultures started with an equal density of 100 mg/L and were cultivated for 16 weeks before dry weight was measured and images were taken. A comparison of the *PpFLOT* mutant lines with a WT control grown with the same density revealed subtle changes, with a discernible color gradient depending on the *PpFLOT* expression level (Figure 3C). This gradient ranged from light green in Δ*PpFLOT-1* to dark brown-green in *PpFLOT*-OEX3 (Figure 3C), and close examination of single protonema cells indicated that increased *PpFLOT* expression led to the development of small round cells in addition to the characteristic cell filaments. These cells resemble vegetative diaspores or brachycytes observed in *P. patens* under exogenous ABA treatments or ABA-mediated stress responses (Arif et al., 2019). These brachycytes are released from the protonema cell network by transforming the surrounding cells into empty tmema cells (Arif et al., 2019), which could not be detected in the *PpFLOT*-OEX lines. These round cells, in contrast to these diaspores, turned reddish-brown with increasing *PpFLOT* levels, contributing to the observed change in culture coloration, especially in the strongest *PpFLOT*-OEX lines. Notably, this coloration expands to the surrounding cell filaments as well (Figure 3C). Interestingly, when *PpFLOT*-OEX protonema was plated on solid media, these structures receded, and after two weeks, they mostly recovered, developing WT-typical gametophores (Figure 3C). Additionally, mature gametophores submerged in liquid media without tissue disruption showed no changes in pigmentation (Supplementary Figure 2).

### Salt and phytohormone sensitivity in *P. patens* are modulated by *PpFLOT* expression levels

To evaluate the responses of the generated mutant lines to elevated salt concentrations, increased osmotic pressure, and exogenous phytohormone treatment, protonema cultures of an equal density (100 mg/L dry weight) were spotted on standard solid medium supplemented with 250 mM NaCl, 300 mM NaCl, 700 mM mannitol, 10 µM 2-*cis*,4-*trans*-abscisic acid (ABA), 10 µM of the auxin analog 1-naphthylacetic acid (NAA) and 10 µM of the cytokinin derivate 6-y-y-(dimethylallylamino)-purine (2-ip). The spotted colonies were cultivated for eight weeks. Under control conditions and 250 mM NaCl treatment, Δ*PpFLOT-1* and *PpFLOT*-OEX lines showed no significant changes in development compared to the WT (Figure 4). However, at a salt concentration of 300 mM, the *PpFLOT*-OEX lines were unable to form colonies whereas both Δ*PpFLOT-1* and the WT exhibited colony formation (Figure 4). To assess the impact of increased osmotic pressure on the *PpFLOT*-OEX lines, which exhibit reduced salt tolerance, protonema cultures of all lines were placed on plates supplemented with 700 mM mannitol. This concentration was chosen to simulate the osmotic pressure equivalent to 300 mM NaCl for *P. patens*, following observations described by Saavedra et al. (2006). All tested lines were able to develop colonies and exhibited growth similar to the WT. However, consistent with our observations in liquid culture, a gradient of pigmentation was observed in the colonies, shifting to a green-brownish hue with increased *PpFLOT* expression (Figure 4). The findings suggest that increased expression of *PpFLOT* is detrimental to the salt tolerance of *P. patens* implicating that PpFLOT most likely acts as a negative regulator in abiotic stress response, either directly or indirectly. Notably, qRT-PCR measurements of the *PpFLOT* expression in WT protonema cultures treated with 250 mM NaCl over 24 h showed a decrease in the expression levels (Figure 5A) at 8 and 24 h. However, this decrease in expression was not statistically significant (ANOVA, p = 0.075) due to a high variance among the biological replicates. Conversely, the expression levels in the salt-treated *PpFLOT*-OEX1 line did not exhibit significant decrease (ANOVA, p = 0.26) (Figure 5A).

**Figure 4:**
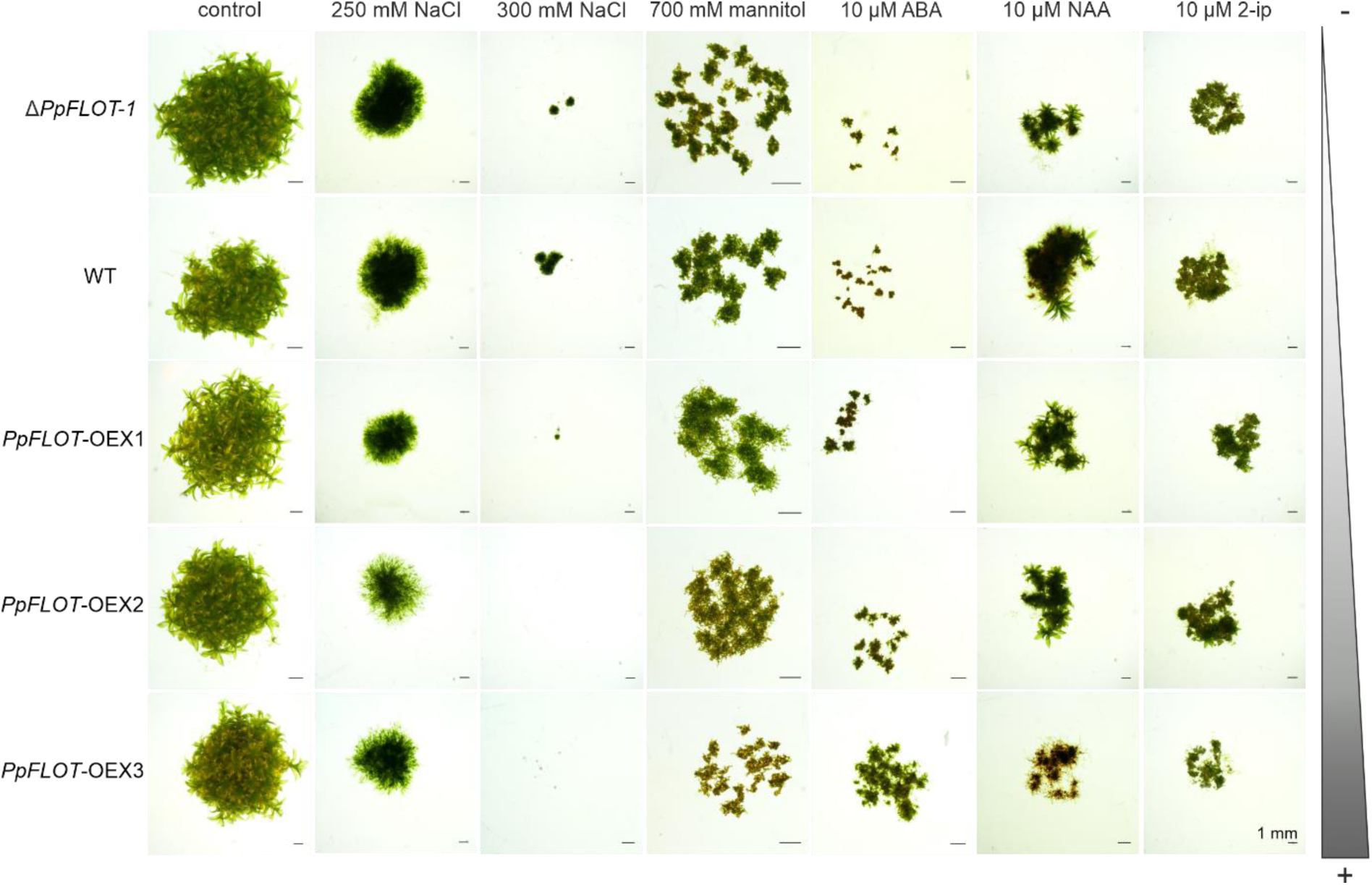
Phenotypic analysis of Δ*PpFLOT-1* and all *PpFLOT*-OEX lines. Δ*PpFLOT-1*, WT, *PpFLOT*-OEX1, *PpFLOT*-OEX2 and *PpFLOT*-OEX3 were inoculated with an equal density (100 mg/L dry weight) on standard solid medium (control) and media supplemented with 250 mM NaCl, 300 mM NaCl, 700 mM mannitol, 10 µM ABA, 10 µM NAA and 10 µM 2-ip and grown for at 8 weeks. The scale bar in all images represents 1 mm. The strength of *PpFLOT* expression is indicated on the right.

**Figure 5:**
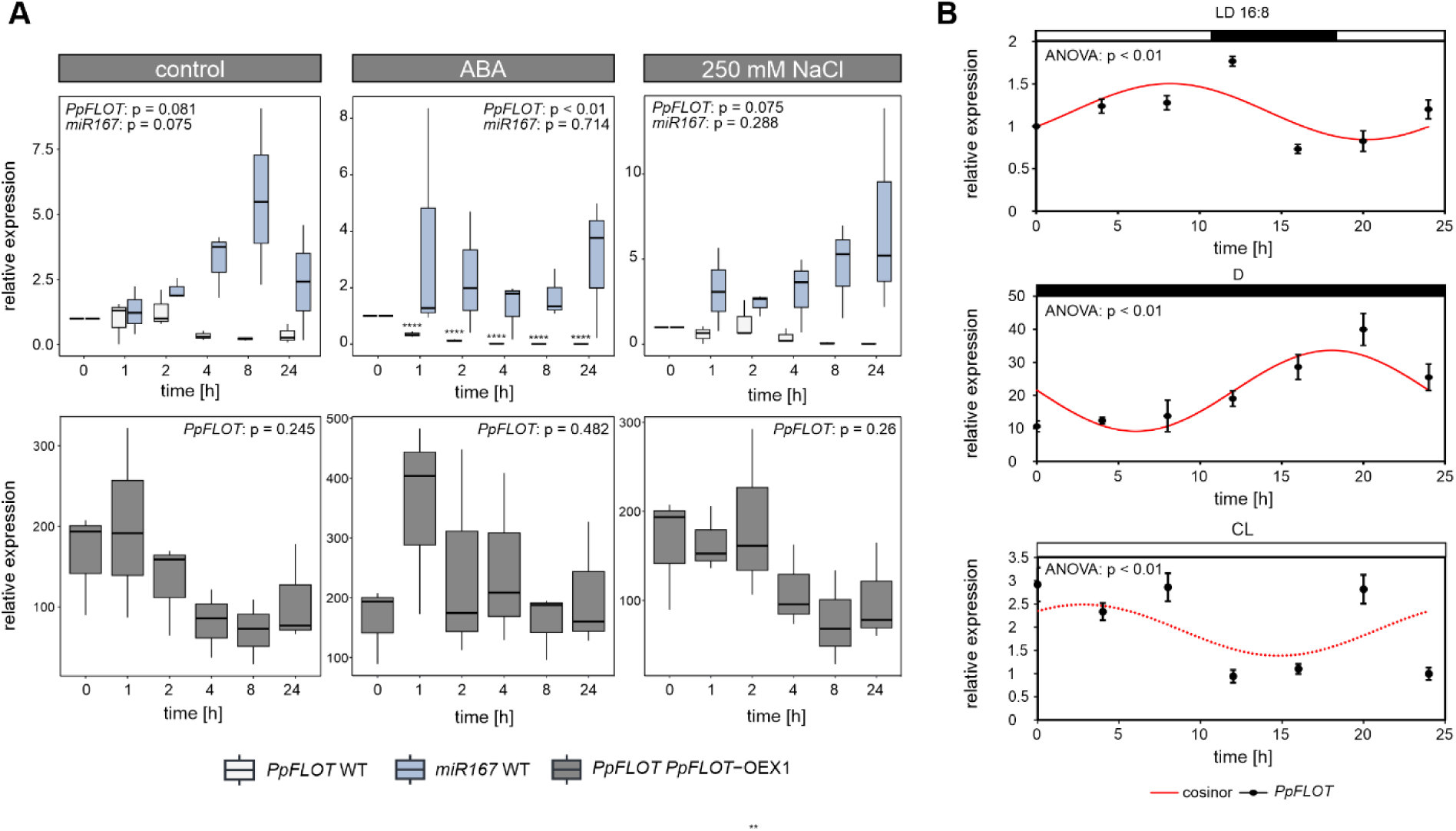
*PpFLOT* expression is regulated by daytime, light, ABA and salt. (A) Depiction of relative gene expression of *PpFLOT* and *miR167* in *P. patens* WT protonema (upper panel) and expression of *PpFLOT* in *PpFLOT*-OEX1 (lower panel) measured over 24 h under control conditions and in response to 10 µM ABA and 250 mM NaCl in triplicates. Analysis performed according to Schmittgen and Livak (2008). ANOVA analysis was conducted to identify statistically significant differences between time points, with p-values provided in the respective boxes. Asterisks denote time points with statistically significant changes in expression identified by Tukey’s HSD test compared to 0 h treatment. * p < 0.05, ** p < 0.01, **** p < 0.0001. (B) Relative gene expression of *PpFLOT* in *P. patens* WT protonema observed over 24 h under long day conditions (LD 16:8), complete darkness (D) and continuous light after adapting for three days to the respective light conditions. Mean values of biological triplicates are depicted with ± SEM (n = 3). Bars above the graphs indicate light conditions at the respective time points (black = light off, white = light on). Light intensity in all conditions was 85–100 µmol/m^2^s. P-values of the ANOVA analyses are given in the respective graphs. Cosinor curve calculated from acrophase, meseor, and amplitude is indicated by a red line. A continuous line indicates statistically significant rhythmicity (p < 0.05) while an interrupted line indicates non-significant rhythmicity (p > 0,05).

During ABA treatment of the respective lines, we further observed improved ABA sensitivity correlating with *PpFLOT* expression, as evidenced by larger *PpFLOT*-OEX3 colonies compared to the Δ*PpFLOT-1* or the WT control (Figure 4). Interestingly, 10 µM ABA treatment of WT protonema cultures led to a statistically significant decrease in *PpFLOT* transcript levels over time (ANOVA, p < 0.01) (Figure 5A). Starting from 1 h after ABA treatment, *PpFLOT* expression significantly decreased and continued to decrease steadily, reaching a relative expression of 0.01 after 24 h (Figure 5A). Treatment with NAA or 2-ip did not result in phenotypic changes for Δ*PpFLOT-1*, *PpFLOT*-OEX1, or *PpFLOT*-OEX2 compared to the treated WT control (Figure 4). In both cases the lines exhibited a similar growth pattern as the wild type, except line *PpFLOT*-OEX3. Upon NAA treatment this line failed to develop gametophores and formed reddish colonies. When treated with 2-ip *PpFLOT*-OEX3 exhibited slightly reduced growth compared to the other lines. (Figure 4). Overall, the observed phenotypic alterations in response to all treatments were correlated with the levels of *PpFLOT* expression.

### The *PpFLOT* transcript is regulated by multiple pathways

The phenotypic analysis of the generated *PpFLOT* mutant lines suggests a putative regulatory function for PpFLOT during abiotic stress responses in *P. patens*. Furthermore, the expression of WT *PpFLOT* is suppressed in response to ABA and salt treatment. Examination of the *PpFLOT* 5’UTR and 1.5 kb the genomic sequence upstream genomic sequence for *cis*-acting regulatory elements using PlantPAN 4.0 (Chow et al., 2019) identified potential binding sites for various transcription factors, APETALA 2 (AP2), basic HELIX-LOOP-HELIX (bHLH), CG-1 DNA-binding domain containing transcription factors (CG-1), ETHYLENE INSENSITIVE 3 (EIN3), GATA transcription factors, DNA binding with ONE FINGER (DOF), LATERAL ORGAN BOUND (LOB) domain transcription factors, V-MYB MYELOBLASTOSIS VIRAL ONCOGENE HOMOLOG (MYB)/ SWITCH_DEFECTIVE PROTEIN 2 (SWI3), ADAPTOR 2 (ADA2), NUCLEAR RECEPTOR COREPRESSOR (N-CoR, TRANSCRIPTION FACTOR III B (TF) (SANT), NO APICAL MERISTEM/ARABIDOPSIS THALIANA ACTIVATING FACTOR/CUP-SHAPED COTYPEDON (NAC), WRKY, MINI CHROMOSOME MAINTENANCE1/AGAMOUS/DEFICIENCE/SERUM RESPONSE FACTOR (MADS) box, BASIC REGION/LEUCINE ZIPPER (bZIP), TATA BINDING PROTEIN (TBP) transcription factors as well as C2H2, TCR, homeodomain and AT-hook motifs. The diverse array of identified putative transcription factor binding sites suggests the involvement of PpFLOT in various biological processes. Interestingly bHLH, NAC, and WRKY, as well as, in some instances, MYB, AP2, DOF, and bZIP transcription factors, are recognized regulators of the abiotic stress response or are themselves regulated by ABA (Golldack et al., 2011; Mizoi et al., 2012; Ambawat et al., 2013; Phukan et al., 2016; Das et al., 2019; Zou and Sun, 2023). Meanwhile, the presence of CG-1 and GATA binding sites suggests potential circadian or light-dependent regulation (da Costa e Silva, 1994; Reyes et al., 2004). Previous studies in rice indicate that AT-hook motifs may interact with light-sensitive phytochromes (Jorge Nieto-Sotelo, 1994). Subsequently, we investigated *PpFLOT* expression for light-dependent or circadian regulation. WT protonema cultures were analyzed for putative circadian regulation by measuring *PpFLOT* transcript levels every 4 h over 24 h. The experiment was conducted under standard growth conditions (16 h light: 8 h dark; LD), complete darkness (D), and continuous light (CL), with all samples pre-adapted to their respective photoperiods for 3 days. ANOVA analysis of the *PpFLOT* expression over 24 h revealed significant changes (p < 0.01) in LD, D and CL cultivated samples. LD samples showed a peak expression at time point (TP) 12, with the lowest *PpFLOT* expression at TP 16, while D samples peaked at TP 20, and TP 0 showed the lowest expression (Figure 5B). In contrast to the other two photoperiod conditions, CL did not display a single maximum or minimum of the expression (Figure 5B). While the overall expression levels of *PpFLOT* in LD and CL samples oscillated between a normalized relative expression of 0.5 – 3, D samples exhibited a higher overall expression level, fluctuating between a normalized relative expression of 10 – 50 (Figure 5B). Intriguingly, the highest *PpFLOT* expression in the LD samples coincided with the beginning of the dark period (Figure 5B), suggesting that *PpFLOT* expression is generally enhanced in the absence of irradiation. To assess if the observed oscillations follow circadian rhythmicity, we conducted a cosinor-based rhythometry analysis (Cornelissen, 2014) for all three groups. Both LD and D expression levels could be fitted to a cosinor curve, meeting the criteria for circadian rhythmicity with p < 0.01. However, the CL group did not fulfill these conditions (p = 0.205). The detected arrhythmicity under CL is likely attributed to a dysfunctionality of the *P. patens* circadian clock at CL conditions, as proposed and demonstrated by Okada et al. (2009) for clock-related genes in *P. patens* (Ichikawa et al., 2004; Ichikawa et al., 2008; Okada et al., 2009; Petersen et al., 2022). Due to the overall higher expression levels, we observed a shift in amplitude and acrophase between the LD and NL samples (Figure 5B). Additionally, a phase delay in D samples compared to LD samples was likely caused by the absence of irradiation impulses (Fukuda et al., 2013). It is noteworthy that in mammals, stress and changes in hormone levels can induce a shift in the expression of genes regulated by the circadian clock (Ota et al., 2021). Therefore, during stress, a delay in the circadian clock of *P. patens* could occur due to increased expression of *PpFLOT* or alterations in phytohormone profiles throughout the day.

In our previous study, altered miRNA biogenesis led to an increase in *PpFLOT* transcript levels (Arif et al., 2022). Subsequently, we searched the genomic *PpFLOT* sequence for a putative miRNA binding site with psRNATarget (Dai et al., 2018), identifying miR167 as the best match with an expectation score of 3.5 and a putative cleavage site in the third exon of the *PpFLOT* sequence located at bp 1898-1918. Relative miR167 expression detected by stem-loop PCR revealed anticorrelating expression between miR167 and *PpFLOT* in WT protonema under control conditions and in response to salt treatment (Figure 5A). However, statistical analysis did not confirm the detected changes in miRNA expression as significant, likely due to high variance between replicates. While these changes in expression were not statistically significant, they suggest a potential regulation of *PpFLOT* on the transcript level. Overall, we identified four potential regulatory mechanisms controlling the *PpFLOT* transcript levels in the *P. patens* protonema cell. The evidence suggests that the expression of *PpFLOT* can be suppressed by ABA-responsive transcription factors. Additionally, *PpFLOT* is circadian regulated, with increased expression when cultivated without light impulses. Moreover, it cannot be dismissed that miR167 may regulate the *PpFLOT* transcript, although further experiments are necessary to confirm this type of regulation. We propose that PpFLOT, located in the chloroplasts of *P. patens*, serves as a scaffolding protein in various biological processes, particularly during the response to abiotic stress and in processes related to the day-night transition of chloroplasts. The diverse methods identified to regulate *PpFLOT* levels transcriptionally or posttranscriptionally in chloroplasts highlight its importance in multiple cellular functions.

### Detection of slight ROS accumulation in *PpFLOT*-OEX lines

All generated *PpFLOT*-OEX lines, displaying lower salt tolerance than the WT or Δ*PpFLOT-1*, potentially attributed to increased ROS accumulation (Pandey et al., 2017) underwent 3,3-diamonobenzidine staining to detect H_2_O_2_ overaccumulation. Protonema samples were incubated for 18 h in the staining solution under CL photoperiod conditions, and subsequent microscopic examination revealed higher H_2_O_2_ levels in *PpFLOT*-OEX lines compared to the WT and Δ*PpFLOT-1* (Figure 6A). Notably, *PpFLOT*-OEX2 and *PpFLOT*-OEX3 exhibited cells with persistent pigmentation, suggesting potential challenges in destaining by boiling in ethanol, making it challenging to ascertain the origin of H_2_O_2_ overaccumulation in these structures. However, a general comparison of stained normal-grown protonema cells across all *PpFLOT*-OEX lines and the WT demonstrated a correlation between *PpFLOT* expression levels and H_2_O_2_ accumulation (Figure 6A).

**Figure 6:**
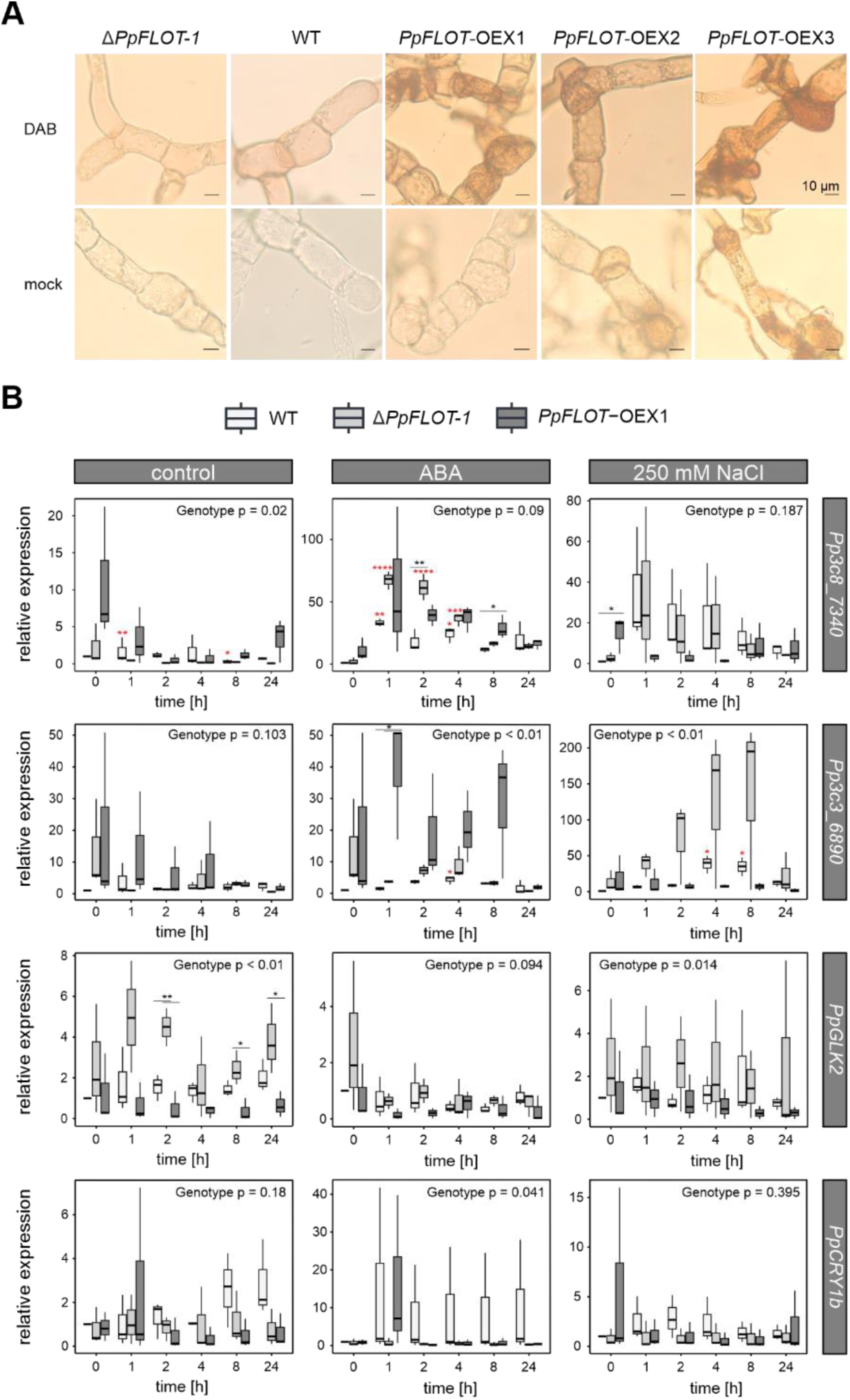
Changes in H_2_O_2_ levels and gene expression corresponding to changes in *PpFLOT* expression. (A) H_2_O_2_ levels were visualized in Δ*PpFLOT-1*, WT, and all three *PpFLOT*-OEX lines through staining with 3, 3-diamonobenzidine. The upper panel depicts images of treated protonema cells, while the lower panel exhibits images of untreated mock controls. The scale bar in all images represents 10 µm. (B) Box plots present qRT-PCR results of relative gene expression for (top to bottom) AP2/ERF domain transcription factor *Pp3c8_7340*, copper transport HMA domain protein *Pp3c3_6890*, *PpGLK2* and *PpCRY1b*, comparing expression levels to WT at 0 h of treatment and normalized against *PpEF1α* following the method by Schmittgen and Livak (2008). The depicted relative gene expression of the respective genes in Δ*PpFLOT-1*, WT and *PpFLOT*-OEX1 protonema is shown under control conditions (left), upon ABA (middle) and salt-(right) treatment measured over 24 h in triplicates. ANOVA was used to determine statistically significant changes in expression between the three genotypes, with p-values indicated in the respective graphs (black asterisk). Tukey’s HSD results for time-dependent expression in one line are marked by red asterisk when significant compared to 0 h of treatment. * p < 0.05, ** p < 0.01, *** p < 0.001, **** p < 0.0001

Elevated H_2_O_2_ levels, indicative of stress in *P. patens*, prompted an examination of transcript levels of abiotic stress-induced genes (Boursiac et al., 2005; Saavedra et al., 2006; Frank et al., 2007; Hauser et al., 2011; Li et al., 2011; Li et al., 2012). These measurements were performed under control conditions and when treated with either 250 mM NaCl or 10µM ABA in WT, Δ*PpFLOT-1* all *PpFLOT*-OEX1. Interestingly, the expression of *AQUAPORINE TIP* (*Pp1s44_31/ Pp3c20_15350*), *9′-*cis *EPOXYCAROTENOID DIOXIGENASE* (*NCED*) (*Pp1s412_7/ Pp3c25_4810*), *TRANSLOCATOR PROTEIN 1* (*TSPO1*) (*Pp1s281_123/ Pp3c2_17540*) and *DEHYDRIN B* (*DHNB*) (*Pp1s442_22/Pp3c5_11880*) barely showed any significant changes in the gene expression among the tested genotypes (Supplementary Figure 3). Specifically, only *PpTSPO1* displayed slightly lower expression in *PpFLOT*-OEX1 compared to the WT and Δ*PpFLOT-1* (ANOVA p = 0.04) in response to salt treatment (Supplementary Figure 3). Conversely, the expression of a mRNA encoding an AP2/ERF domain transcription factor *Pp1s8_127/ Pp3c8_7340* changed significantly between the genotypes under control conditions (ANOVA, p = 0.02) since *Pp3c8_7340* (AP2/ERF) exhibited significantly (Tukey’s HSD, p = 0.0235) higher expression in *PpFLOT*-OEX1 compared to Δ*PpFLOT-1*. Significant upregulation of *Pp3c8_7340* (AP2/ERF) was observed at 0 h of salt treatment (Tukey’s HSD, p = 0.0429) and at 8 h of ABA treatment (Tukey’s HSD, p = 0.0229) in *PpFLOT*-OEX1 compared to the WT. Notably, both the WT and Δ*PpFLOT-1* samples treated with ABA for 1, 2, and 4 h exhibited a significant increase in *Pp3c8_7340* (AP2/ERF) expression compared to untreated samples (ANOVA, p < 0.01). Statistically significant differences in expression levels between the genotypes were observed at 2 h (Tukey’s HSD Δ*PpFLOT-1* vs WT, p < 0.01) and 8 h (Tukey’s HSD *PpFLOT*-OEX1 vs WT, p = 0.0295). However, the overall changes in expression between the tested genotypes in response to ABA treatment were not statistically significant (ANOVA, p = 0.09) (Figure 6B). Similarly, upon salt treatment, the overall change in expression between the tested genotypes was not significant (ANOVA, p = 0.187). At 0 h, a significantly higher expression of *Pp3c8_7340* (AP2/ERF) was detected in *PpFLOT*-OEX1 (Tukey’s HSD *PpFLOT*-OEX1 vs WT, p = 0.0429) (Figure 6B). This upregulation in untreated samples of *PpFLOT*-OEX1 is most likely an attempt to suppress the ectopic *PpFLOT* expression. Heavy metal-associated domain (HMA) containing proteins are vital for heavy metal transport, detoxification in plants, and respond to various stresses (Li et al., 2020; Barr et al., 2023). Proteins in this family are upregulated in response to drought, cold, hypoxia, and bacterial infection (Barr et al., 2023). The gene expression of the copper transport protein Pp1s296_27/Pp3c3_6890, encoding such an HMA domain, exhibited significant differences in expression between the three genotypes upon ABA and salt treatment (ANOVA, p < 0.01). For instance, while all genotypes showed increasing *Pp3c3_6890* levels in response to ABA, this upregulation was extreme in *PpFLOT*-OEX1 1 h after ABA treatment compared to the other two lines (Tukey’s HSD, p < 0.05) (Figure 6B). On the other hand, in response to salt Δ*PpFLOT-1* displayed an extreme upregulation of *Pp3c3_6890* (Cu-HMA) compared to both WT (Tukey’s HSD, p < 0.01) and *PpFLOT-*OEX1 (Tukey’s HSD, p < 0.01). Comparatively *Pp3c3_6890* (Cu-HMA) expression is subdued in *PpFLOT*-OEX1 in response to salt (Figure 6B). Despite the salt-sensitive phenotype of *PpFLOT*-OEX lines, only minor changes in the expression levels of salt and ABA-induced genes were detected upon induction. For instance, *Aquaporine TIP* exhibited slight repression in *PpFLOT*-OEX1 at all TP compared to 0 h under control conditions. Comparatively, *PpFLOT*-OEX1 showed a peak in *Aquaporine TIP* expression at 2 h instead of 24 h (Supplementary Figure 3). In the WT, *NCED* expression significantly increased in response to ABA (ANOVA, p = 0.015) and salt treatment (ANOVA, p < 0.01). While this change in expression was also detected in *PpFLOT*-OEX1 and Δ*PpFLOT-1*, it was not statistically significant (*PpFLOT*-OEX1: ANOVA salt p = 0.585, ABA p = 0.427 and Δ*PpFLOT-*1: ANOVA salt p = 0.629; ABA p = 0.078) (Supplementary Figure 3). Similarly, no changes in the expression of *PpDHNB* were detected among the three genotypes (ANOVA, p = 0.865) over 24 h. However, at 24 h, *PpDHNB* expression was significantly upregulated in the WT compared to Δ*PpFLOT-1* and *PpFLOT*-OEX1 (Tukey’s HSD, p < 0.01). Additionally, the peak expression of *PpDHNB* upon salt treatment for both WT and *PpFLOT*-OEX1 occurred at 2 h, while in Δ*PpFLOT-1*, it peaked at 24 h (Supplementary Figure 3). Most abiotic stress markers behaved similarly to the WT, suggesting no upregulation of ABA biosynthesis or other known abiotic stress-sensing pathways in in Δ*PpFLOT-1* and *PpFLOT*-OEX lines.

Abnormal expression of stress-induced genes and increased H_2_O_2_ levels may impact retrograde signaling pathways, involving chloroplast-to-nucleus communication using ROS as signaling molecules (Locato et al., 2018; Li and Kim, 2022). The gene *GOLDEN 2-LIKE PROTEIN 2* (*Pp3c11_21140*) (*GLK2*), associated with retrograde signaling (Sun et al., 2022; Zeng et al., 2023), exhibited significant expression changes under control conditions and salt treatment in WT, Δ*PpFLOT-1* and all *PpFLOT*-OEX lines (ANOVA, salt p = 0.014, cont p < 0.01). Δ*PpFLOT-1* displayed consistently higher *PpGLK2* expression than WT and *PpFLOT*-OEX1 under control conditions (Tukey’s HSD, p <0.01) (Figure 6B). In response to salt treatment, *PpFLOT*-OEX1 showed significantly lower *PpGLK2* expression (Tukey’s HSD, p < 0.0119) compared to Δ*PpFLOT-1* but not compared to the WT (Tukey’s HSD, p = 0.581) (Figure 6B). Another retrograde signaling gene, *CRYPTOCHROME 1b* (*Pp3c7_20480V3*) (*PpCRY1b*), displayed a statistically significant expression change (ANOVA, p = 0.041) between the genotypes upon ABA treatment. *PpCRY1b* was generally expressed at higher levels in WT than Δ*PpFLOT-1* (Tukey’s HSD, p < 0.0356) except at 1 h. However, under control conditions, *PpCRY1b* showed higher transcript levels at 8 and 24 h after start of the measurements compared to Δ*PpFLOT-1* and *PpFLOT*-OEX1, with no statistically significant differences under control conditions (Figure 6B).

### Increased chlorophyll a/b content correlates with strength of *PpFLOT* expression

The total chlorophyll content of WT, Δ*PpFLOT-1*, *PpFLOT*-OEX1, *PpFLOT*-OEX2, and *PpFLOT*-OEX3 was assessed to determine if the observed *PpFLOT*-dependent change in pigmentation resulted in altered chlorophyll content in the *PpFLOT* mutant lines. Chlorophyll a and b were extracted using 80 % acetone from equal amounts of protonema harvested per line. After separating cell debris, chlorophyll absorption was measured, and total chlorophyll content was calculated following the method outlined by Frank et al. (2005). Significant changes (ANOVA, p < 0.01) in the chlorophyll content were observed between *PpFLOT* mutant lines and the WT control. Δ*PpFLOT-1* exhibited only half as much chlorophyll as the WT, with 0.4 mg chl/g dry weight (Figure 7A). Surprisingly, the *PpFLOT*-OEX1 line exhibited a similar amount of chlorophyll (0.49 mg chl/g dry weight). In contrast, both *PpFLOT*-OEX2 and *PpFLOT*-OEX3 had significantly higher chlorophyll content, with 2.6 and 2.5 mg chl/g dry weight, respectively (Figure 7A). This suggests an enhanced chlorophyll biosynthesis, potentially leading to increased photosynthetic activity. To assess this, we performed Pulse-Amplitude-Modulation (PAM) measurements to quantify the chlorophyll fluorescence and calculate the quantum yield of the photosystem II (PSII) activity and non-photochemical quenching during photosynthesis activation by light. As previously mentioned, we observed massive differences in the pigmentation between *PpFLOT* mutant lines grown in liquid cultures and those cultivated on solid media. Therefore, we conducted separate measurements for mature gametophores grown on solid medium and protonema liquid cultures with the same number of biological replicates (n = 9) using identical parameters. Interestingly, while statistical analysis of PSII activity among all tested genotypes in the gametophores revealed significant differences (ANOVA, p = 0.023), only *PpFLOT*-OEX1 displayed a lower PSII activity compared to WT and Δ*PpFLOT-1*, while *PpFLOT*-OEX2 and 3 showed no differences in activity compared to the WT. During non-photochemical quenching, we found no changes between the WT, Δ*PpFLOT-1*, and all *PpFLOT*-OEX lines in the gametophores (ANOVA, p =0.34) (Figure 7B). To ensure comparability between the gametophore and the protonema samples, we measured fluorescence at 450 nm and an actinic light intensity of 55 µmol/m^2^s photosynthetically active radiation (PAR). However, we were unable to detect the fluorescence of *PpFLOT*-OEX3 protonema under these conditions, and the measuring light parameter had to be adjusted to an intensity of 60 µmol/m^2^s PAR to detect a fluorescence signal. Therefore, PAM measurements of *PpFLOT*-OEX3 protonema were excluded from the analysis to maintain comparability between all measurements. In contrast to the fluorescence measurements in gametophores, we detected statistically significant differences in the Y(II) and NPQ/4 (ANOVA, p < 0.01) between Δ*PpFLOT-1* and *PpFLOT*-OEX1 and 2 protonema cultures. Compared to the WT samples, neither the Δ*PpFLOT-1* nor the two *PpFLOT*-OEX lines showed any significant differences in the calculated photosynthetic activity (Tukey’s HSD, Δ*PpFLOT-1* vs WT p = 0.898, *PpFLOT*-OEX1 vs WT p = 0.0719, *PpFLOT*-OEX1 vs WT p = 0.165). However, a comparison between the Δ*PpFLOT-1* and the *PpFLOT*-OEX lines revealed a significantly higher quantum yield of PSII in the *PpFLOT*-OEX lines (Tukey’s HSD, *PpFLOT*-OEX1 vs Δ*PpFLOT-1* p < 0.01, *PpFLOT*-OEX2 vs Δ*PpFLOT-1* p = 0.0298) compared to the knockout line (Figure 7B). This change in activity was also detected during non-photochemical quenching. For instance, both *PpFLOT*-OEX lines process less energy by non-photochemical quenching than Δ*PpFLOT-1* (ANOVA, p < 0.01; Tukey’s HSD, *PpFLOT*-OEX1/2 vs Δ*PpFLOT-1* p < 0.01) (Figure 7B). We found that the level of photosynthetic activity in protonema cultures slightly increases in correlation with the increased expression of *PpFLOT*. However, a statistical comparison between the WT and all *PpFLOT* mutant lines did not yield any significant results. Interestingly, we detected a significantly higher (ANOVA, p < 0,01) NPQ/4 signal in Δ*PpFLOT-1* compared to *PpFLOT*-OEX1 and 2 (Tukey’s HSD, p < 0.01). The absence of PpFLOT in the knockout line results in lower photosynthetic activity and increased non-photochemical quenching, potentially due to reduced chlorophyll levels. We suggest that adjusting the amount of *PpFLOT* in chloroplasts of protonema cells can fine-tune the photosynthetic activity to accommodate the specific needs of the respective cells. This seems to be relevant only during the protonema life stage of *P. patens* as *PpFLOT* overexpression, with the exception of *PpFLOT*-OEX1, has barely an effect on the photosynthetic activity of the gametophores.

**Figure 7:**
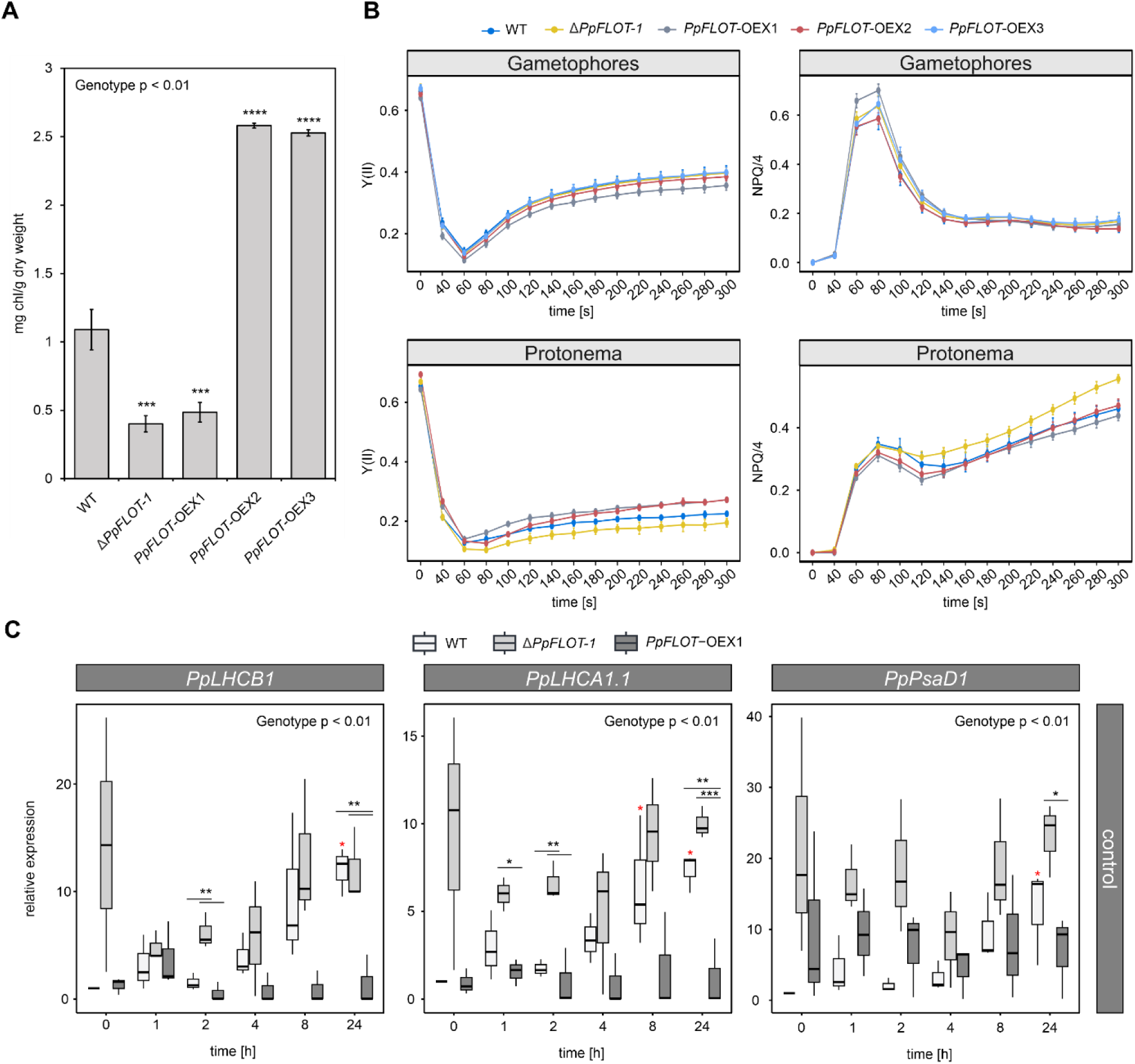
Effects of changes in *PpFLOT* expression on chlorophyll content and photosynthetic activity. (A) Chlorophyll content of 0.4 g protonema from WT, Δ*PpFLOT-1* and all *PpFLOT-OEX* lines. Mean values ± SEM (n =3) are shown. ANOVA results are provided, and statistically significant (p < 0.05) differences in the *PpFLOT* mutants compared to the WT were determined by Tukey’s HSD test and are indicated by asterisks. * p < 0.05, ** p < 0.01, *** p < 0.001, **** p < 0.0001 (B) Effect of *PpFLOT* expression on Y(II) and NPQ/4 in the gametophore and protonema life stage of *P. patens*. Mean values ± SD (n = 9) are shown. Measurements were performed after 3 h of dark adaptations and actinic light of 55 µmol/m^2^s at 450 nm for 315 s after saturation pulse stimulation. (C) qRT-PCR results shown as box blots of relative gene expression of *PpLHCB1* (Pp3c2_35930), *PpLHCA1.1* (Pp3c13_14980), *PpPsaD1* (Pp3c16_23780) over 24 h relative to WT expression at 0 h, normalized against *PpEF1α* as described by Schmittgen and Livak (2008). Depicted is the relative gene expression of the respective genes in Δ*PpFLOT-1*, WT and *PpFLOT-OEX1* protonema under control conditions measured in biological triplicates. Statistically significant changes in expression between the three genotypes were determined by ANOVA, with p-values provided in the respective graphs. Significant differential gene expression at a specific time point is marked by black asterisks. Results of the Tukey’s HSD for the time-dependent expression within the same genotype are marked by red asterisks when significant compared to 0 h of treatment. * p < 0.05, ** p < 0.01, *** p < 0.001, **** p < 0.0001

Since photosynthetic activity is slightly altered in the *PpFLOT* mutant protonema liquid cultures and these lines display changes in the chlorophyll biosynthesis rate, we measured the transcript levels of photosystem one (PSI) and two (PSII) related components. Under control conditions, the expression of the *LIGHT-HARVESTING COMPLEX B1* (*LHCB1*) (*Pp3c2_35930*) displayed significant (ANOVA, p < 0.01) differential expression when compared between WT, Δ*PpFLOT-1* and *PpFLOT*-OEX1 over 24 h. Δ*PpFLOT-1* showed significantly higher *PpLHCB1* expression compared to the WT (Tukey’s HSD, p = 0.0347) and *PpFLOT*-OEX1 (Tukey’s HSD, p < 0.0347). Although not statistically significant, *PpLHCB1* expression appears subdued in the *PpFLOT*-OEX1 line in general (Figure 7C).

Meanwhile, the expression of the *LIGHT-HARVESTING COMPLEX A1.1* (*LHCA1.1*) (*Pp3c13_14980*) not only changes under control conditions (ANOVA, p < 0.01) (Figure 7C), but also in response to salt (ANOVA, p = 0.031) and ABA treatment (ANOVA, p < 0.01) (Supplementary Figure 3). Under all conditions, *PpLHCA1.1* showed a lower expression in the *PpFLOT*-OEX1 line compared to both the WT and Δ*PpFLOT-1* (Figure 7C, Supplementary Figure 3). A similar trend was observed for the expression of the *PSI SUBUNIT D1* (*PsaD1*) (*Pp3c16_23780*). Comparison among the three tested lines revealed a statistically significant change in expression under control conditions (ANOVA, p < 0.01) (Figure 7C), ABA (ANOVA, p = 0.013) and salt treatment (ANOVA, p = 0.019) (Supplementary Figure 3). Under control conditions, *PpPsaD1* expression was significantly higher in the Δ*PpFLOT-1* line compared to the WT (Tukey’s, p < 0.01) and *PpFLOT*-OEX1 (Tukey’s, p < 0.01). Overall, the alterations in chlorophyll content, photosynthetic activity of PSII, and the expression of PS antenna-complex components in correlation with *PpFLOT* expression indicate an involvement of PpFLOT in the photosynthesis of *P. patens*.

### Zeaxanthin level is depleted in correlation to *PpFLOT* expression

The observed color change in the *PpFLOT*-OEX liquid cultures may result from specific alterations in the pigment profile compared to the WT. Thus, the pigment composition of Δ*PpFLOT-1,* all *PpFLOT*-OEX lines, and a WT control was analyzed by liquid chromatography-tandem mass spectrometry (LC-MS) in four replicates. Among 24 detected pigments, 19 could not be identified. To identify differentially accumulated pigments, statistical analysis was conducted using Perseus v 2.0.11 (MaxQuant) (Tyanova et al., 2016). Pigments were classified as DE when both the p-value and the permutation-based false discovery rate (FDR; q-value) of the Student’s *t*-test of Δ*PpFLOT-1* and all *PpFLOT*-OEX lines using the WT as a control group were less than 0.05. The analysis employed the log_2_ transformed fold change (FC) of the relative pigment abundance. It was found that two detected pigments, lutein and unknown pigment 16, were differentially accumulated in all groups. Additionally, zeaxanthin and unknown pigment 10, displayed an altered abundance in the *PpFLOT*-OEX lines (Figure 8A, Supplementary Table 2) that correlates with *PpFLOT* expression. Notably, all shared differentially accumulated pigments of the *PpFLOT* mutant lines were downregulated compared to the WT (Figure 8A, Supplementary Table 2). However, while the detected amount of these pigments decreased in the *PpFLOT* mutant lines, only zeaxanthin and unknown pigment 10 decreased in correlation with the *PpFLOT* expression (Figure 8A). As zeaxanthin is part of the xanthophyll cycle, we further analyzed the accumulation of its other main components alongside lutein in the *PpFLOT* mutant lines (Figure 8B).

**Figure 8:**
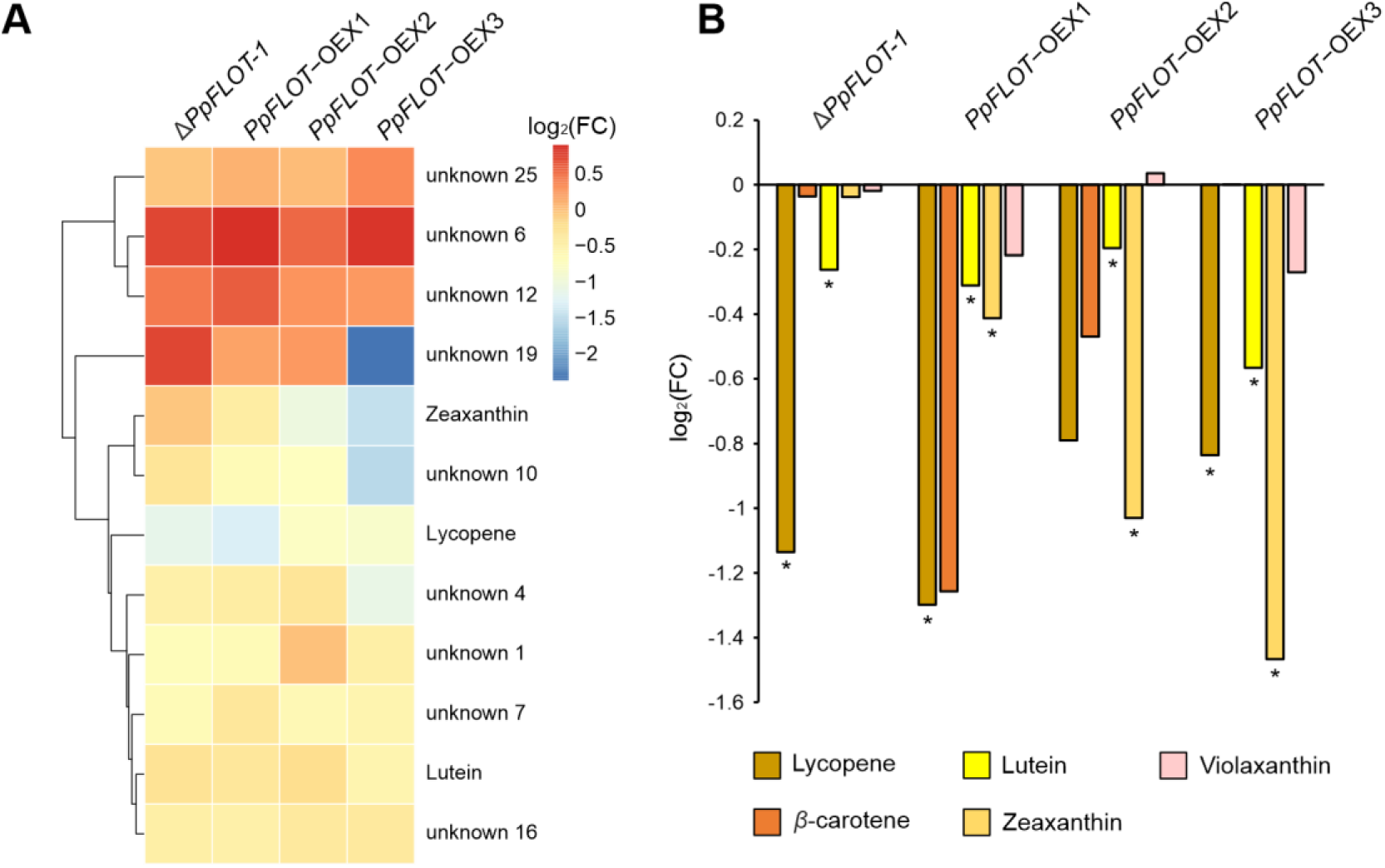
Pigment composition of all *PpFLOT* mutant lines. (A) The clustered heatmap displays log_2_(FC) values of identified pigments, which were deemed differentially accumulated by Student’s *t*-test (p < 0.05) and the permutation-based false discovery rate (FDR) (p < 0.05) in at least one *PpFLOT* mutant line in comparison to the WT. (B) The log_2_(FC) of key xanthophyll cycle components, including lycopene, lutein, violaxanthin, β-carotene, and zeaxanthin is shown for Δ*PpFLOT-1* and all *PpFLOT*-OEX lines compared to the WT. Differentially accumulated pigments identified by Student’s *t*-test and FDR are denoted with an asterisk.

It is noteworthy that the lycopene pool was significantly depleted in Δ*PpFLOT-1*, *PpFLOT*-OEX1 and 3 when compared to the WT (Figure 8A, Supplementary Table 2). β-carotene also showed a slight depletion in *PpFLOT*-OEX1 and 2 but recovered in *PpFLOT*-OEX3. The abundance of violaxanthin exhibited only minor changes in the *PpFLOT* mutant lines compared to the WT (Figure 8B). No other differentially accumulated components of the xanthophyll cycle were detected or identified. Interestingly unknown pigments 6, 12, 19, and 25 all showed increased abundance in at least three *PpFLOT* mutant lines compared to the WT. Identification of these unknown pigments may elucidate the molecular processes responsible for the observed change in culture coloration. It is also interesting to note that with the exception of zeaxanthin, no gradual change in the analyzed pigment levels can be detected since *PpFLOT*-OEX2 seems to behave differently from *PpFLOT*-OEX1 and 3. These differences most likely occurred due to a higher variability between the technical replicates and extreme outliers in this group compared to the other *PpFLOT* mutant lines.

### Changes in the proteome of *PpFLOT*-OEX lines convey influence on metabolic pathways in chloroplasts

To identify potential proteins that could explain the observed phenotype and altered salt tolerance, a proteomics analysis was carried out. Since the change in protonema cell shape and coloration, as well as the altered salt tolerance, was limited to PpFLOT-OEX lines, the proteomic analysis was carried out for the three *PpFLOT*-OEX and a WT control. For this analysis 50 mg fresh weight protonema liquid culture of WT control and all *PpFLOT*-OEX lines were harvested in four replicates and LC-MS was conducted. First analysis of the generated proteomic data confirmed that the increase of *PpFLOT* transcript in all three *PpFLOT*-OEX lines translated into correspondingly higher protein levels. The detected log_2_(FC) of PpFLOT abundance in *PpFLOT*-OEX 1, 2 and 3 lines compared to the WT amounted to 8.12, 8.88 and 9.79, respectively. The statistical analysis of the generated proteomics data was performed with Perseus (Tyanova et al., 2016). To identify statistically significant differentially expressed protein groups (DEP), we performed ANOVA analysis and used the permutation-based FDR of 0.05. A total of 2163 detected protein groups were identified, with 1484 showing significant DE between at least two of the analyzed genotypes (Supplementary Table 3). By visualizing the z-scores of the DEP using a clustered heatmap, a shift in the protein expression correlated with the amount of synthesized PpFLOT (Figure 9A) was shown. Cluster one contained proteins whose expression increased with enhanced PpFLOT expression, while the expression of protein groups in cluster three was suppressed (Figure 9A, Supplementary Table 3). Interestingly, clusters two and four both showed changes in protein abundance in P*pFLOT*-OEX1 and 2 compared to the WT, while the protein expression profile of *PpFLOT*-OEX3 was WT like (Figure 9A, Supplementary Table 3). To identify the molecular functions, cellular components, and biological processes influenced by *PpFLOT* overexpression, a gene ontology (GO) enrichment analysis was performed using shinyGO v 0.8 (Ge et al., 2020). The majority of affected cellular components are part of the thylakoids and the chloroplasts, supporting the localization of *PpFLOT* associated to thylakoid membranes. This implies a function during photosynthetic or photosynthesis-related processes, most likely by assisting protein complex assembly (Figure 9B). Interestingly among the affected GO-terms in the category molecular function were copper ion binding, structural molecular activity, and oxidoreductase activity, particularly focusing on oxidoreductase activity when acting on CH-OH groups (Figure 9B). The detected enriched biological processes also suggest an involvement of PpFLOT in photosynthesis and biosynthesis of carbohydrates, amides, organic acids, and peptides (Figure 9B, Supplementary Tables 3 and 4).

**Figure 9:**
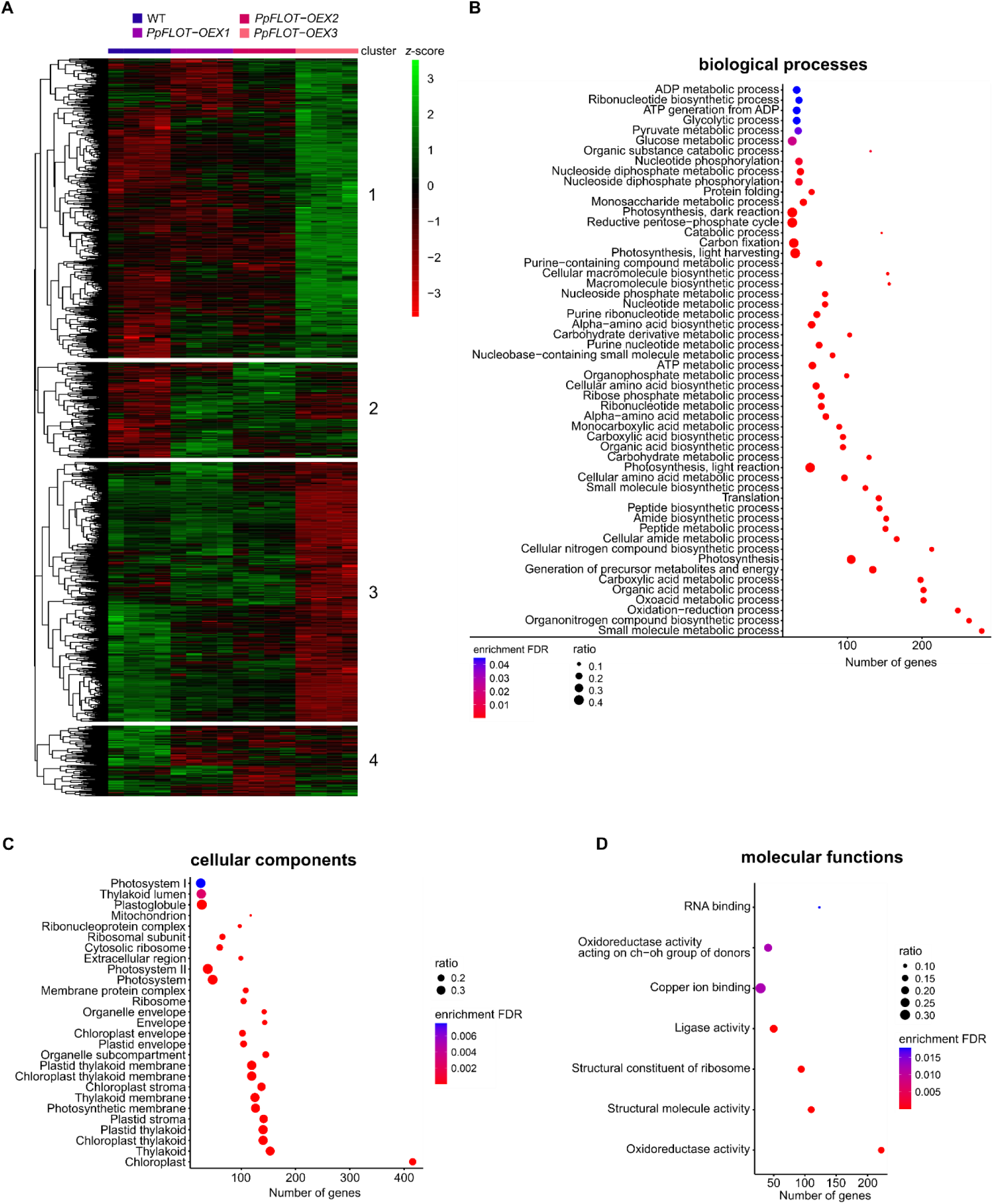
Proteomics and GO term analysis of all *PpFLOT*-OEX lines compared to *P. patens* WT. (A) Clustered heatmap of z-score transformed log_2_(LFQ intensities) values of all differentially expressed protein groups (DEP) identified by ANOVA (p < 0.05) and a significant permutation-based false discovery rate (FDR) (q < 0.05). The depicted protein groups are DEP between at least two of the analyzed genotypes. Clusters are numbered and proteins contained in each cluster are listed in Supplementary Table 3. (B - D) Results of the GO term analysis performed by shinyGO (Ge et al., 2020) and sorted after (B) biological processes, (C) cellular components, and (D) molecular functions. Shown are the significant (p < 0.05) GO terms also displaying a significant enrichment (FDR) q < 0.05. The number of detected proteins and the ratio between total genes of a pathway and the detected genes/proteins of that pathways are given.

To identify DEP in each *PpFLOT*-OEX line compared to the WT specifically, we performed a Student’s *t*-test for each overexpression line. Protein groups with a p-value and a permutation-based FDR of 0.05 and a log_2_(FC) ≤ −1 and ≥ + 1 were identified as DEP (only main IDs shown). This way, we identified 89 DEPs (76 up and 13 down) in *PpFLOT*-OEX1, 92 in *PpFLOT*-OEX2 (62 up and 30 down), and 216 in *PpFLOT*-OEX3 (135 up and 81 down) (Figure 10A, 10B, Supplementary Table 5). The majority of these DEPs are upregulated in all three overexpression lines, and only a fraction of them are downregulated (Figure 10C).

**Figure 10:**
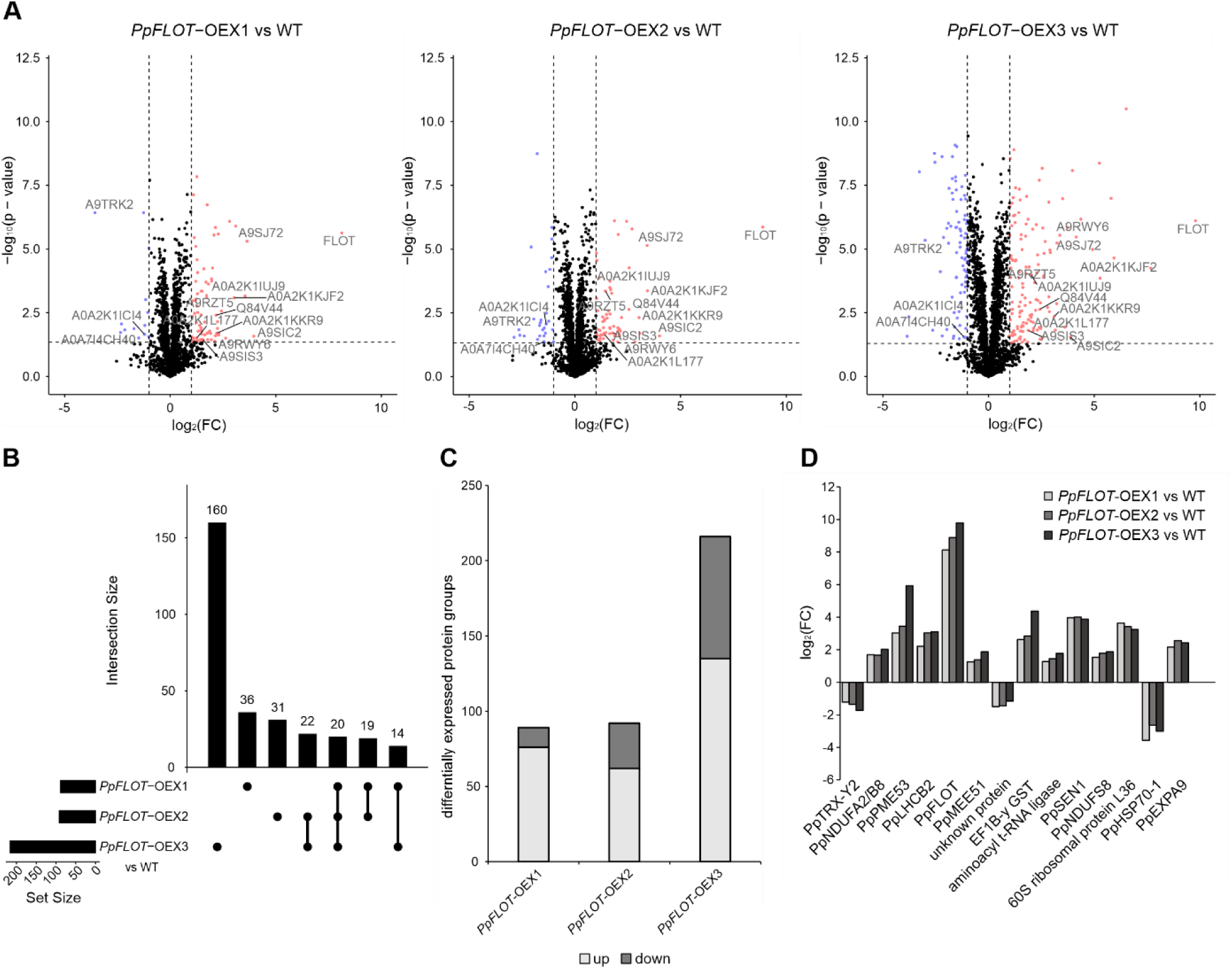
Identification of differentially expressed protein groups in all *PpFLOT*-OEX lines. (A) Volcano plots of all differentially expressed protein groups (DEP) identified by Student’s *t*-test with p < 0.05 and permutation-based false discovery rate (FDR) q < 0.05 using WT expression as the control group. All DEP exhibited a log_2_(FC) of < - 1 and > 1. UniProt protein ID was added to the DEP common in all *PpFLOT*-OEX lines and either correlated with the PpFLOT expression or showed a log_2_(FC) of < - 2 and > 2. Downregulated DEPs are denoted in blue and upregulated DEPs are denoted in red. (B) UpSet plot of all identified DEP in the three *PpFLOT*-OEX lines. (C) Bar chart showing the amount of up- or downregulated DEP in all three *PpFLOT*-OEX lines. (D) Bar chart shows 14 common DEP between the three *PpFLOT*-OEX lines either showing a PpFLOT correlating expression or a log_2_(FC) of < - 2 and > 2. DEP: PpTRX-Y2 (A0A2K1ICI4); PpNDUFA2/B8 (A0A2K1IUJ9); PpPME53 (A0A2K1KJF2); PpLHCB2 (A0A2K1KKR9); PpFLOT (A0A2K1KVH5); PpMEE51 (A0A2K1L177); unknown protein (A0A7I4CH40); EF1B-γ GST (A9RWY6); aminoacyl t-RNA ligase (A9RZT5); PpSEN1 (A9SIC2); PpNDUFS8 (A9SIS3); 60S ribosomal protein L36 (A9SJ72); PpHSP70-1 (A9TRK2); PpEXPA9 (Q84V44).

However, the number of downregulated DEP increases with the strength of *PpFLOT* expression (Figure 10C, Supplementary Table 5). Our analysis shows that *PpFLOT*-OEX3 displays the largest modification in its proteomics profile with 160 DEP, compared to *PpFLOT*-OEX1 and *PpFLOT*-OEX2, which display 36 and 31 DEP, respectively, that are specific to these lines and do not overlap with any of the other two overexpression lines. However, 20 DEP were found to be differentially accumulated in all three lines (Figure 10 B, Supplementary Table 5). Notably, among these, 14 proteins (including PpFLOT) displayed either a log_2_(FC) ≤ −2 and ≥ + 2 or a correlated DE to the strength of *PpFLOT* expression (Figure 10D, Supplementary Table 5). THIOREDOXIN Y2 (PpTRX-Y2) (Pp3c26_10440/A0A2K1ICI4), HEAT SHOCK COGNATE PROTEIN 70-1 (PpHSP70-1) (Pp3c4_21500/ A9TRK2) and one unknown protein (A0A7I4CH40) showed decreased protein levels. TRX-Y2 plays a role in oxidative stress response signaling pathways in chloroplasts (Geigenberger et al., 2017; Wittmann et al., 2021), while PpHSP70-1 is a cytosolic HSP70 involved in protein trafficking and maintenance (Cazalé et al., 2009; Shi and Theg, 2010; Leng et al., 2017). Double mutants of functionally redundant HSP70-1 and −4 in *A. thaliana* exhibited salt hypersensitivity and ABA hyposensitivity (Leng et al., 2017). Conversely, HSP70-1 overexpression lines showed increased tolerance to abiotic stress, including salinity stress (Cazalé et al., 2009). Therefore, the downregulation of both PpTRX-Y2 and PpHSP70-1 might contribute to the altered salt sensitivity and response to enhanced osmotic pressure in *PpFLOT*-OEX lines. Additionally, a homolog to *A. thaliana* PECTIN METHYLESTERASE 53 (PpPME53) (Pp3c5_12660/A0A2K1KJF2) and EXPANSIN A9 (PpEXPA9) (Pp3c12_4560/Q84V44) are both upregulated in all *PpFLOT*-OEX lines. Since both proteins are involved in restructuring the cell wall (Li et al., 2002; Cosgrove, 2015; Gigli-Bisceglia et al., 2022), and cell wall integrity is an important factor during salt sensing (Shin et al., 2021; Gigli-Bisceglia et al., 2022), these differences in expression might influence the salt sensing capability of the *PpFLOT*-OEX lines. Besides these cell wall-related proteins, two proteins related to electron transport in mitochondria, NADH DEHYDROGENASE UBIQUINONE IRON-SULFUR PROTEIN 8 (PpNDUFS8) (Pp3c15_4330/A9SIS3) and NADH-ubiquinone oxidoreductase B8 subunit (PpNDUFA2/B8) (Pp3c20_8510/A0A2K1IUJ9) were upregulated, indicating increased respiration due to *PpFLOT* overexpression (Klodmann and Braun, 2011; Domergue et al., 2022). Also, the *A. thaliana* MATERNAL EFFECT EMBRYO ARREST 51 (MEE51) homolog (Pp3c2_11980/A0A2K1L177) is upregulated. PpMEE51 belongs to the phosphofructokinase family, and its upregulation might indicate an increasingly anoxic environment in the *PpFLOT*-OEX lines since both ATP-dependent phosphofructokinases (PFK) and pyrophosphate-fructose-6-phosphotransferases (PFP) are upregulated in *O. sativa* upon anoxia (Mustroph et al., 2013). Furthermore, knockout of *PpNDUFS8* homolog in *A. thaliana* led to increased lipid content, suggesting that its higher expression in *P. patens* might also lead to changes in the lipid profile (Domergue et al., 2022). An EF1B-γ glutathione S-transferase (GST) (Pp3c18_20060/A9RWY6) is also upregulated. In plants, GSTs are not only involved in the abiotic but also in the biotic stress response (Gullner et al., 2018; Hernández Estévez and Rodríguez Hernández, 2020). Interestingly, while our transcript expression analysis in *PpFLOT*-OEX1 revealed suppression of *PpLHCB1* and *PpLHCA1.1*, we detected increased levels of PpLHCB2 (Pp3c5_22920/A0A2K1KKR9) in all three *PpFLOT*-OEX lines. It is noteworthy that PpLHCB2 was the sole PS antenna component that displayed a change in protein level. Additionally, a Rhodanese-containing protein, homologous to *A. thaliana* SENESCENCE 1 (AtSEN1) also known as DARK INDUCIBLE 1 (AtDIN1) (Pp3c16_18500/A9SIC2), exhibited elevated protein levels. Similar to *PpFLOT*, *AtSEN1* expression is induced in darkness and exhibits increased transcript levels in response to bacterial infection (Schenk et al., 2005; Fernandez-Calvino et al., 2016). Furthermore, *AtSEN1* participates in senescence (Fernandez-Calvino et al., 2016). Thus, the dependence of PpSEN1 accumulation on PpFLOT suggests a role for PpFLOT in the pathogen response.

### Overexpression of *PpFLOT* leads to changes in the fatty acid profile an increases monogalactosyldiacylglycerol (MGDG) accumulation

Chloroplasts do not only serve as the site of photosynthesis, but also for multiple metabolic pathways, including fatty acid (FA) biosynthesis (Block et al., 2007; Eberhard et al., 2008; Johnson, 2016; He et al., 2020). Fatty acids are essential building blocks for many metabolic compounds, including the cuticle layer that covers the outer cell wall of *P. patens* and plasma membrane components (Renault et al., 2017; Resemann et al., 2019; Batsale et al., 2021). Previous studies have shown that plasma membrane fluidity and cell wall structure are crucial factors during the abiotic stress tolerance, particularly during drought and cold stress of *P. patens* and *A. thaliana* (Bhyan et al., 2012; Barrero-Sicilia et al., 2017; Batsale et al., 2021). Moreover, increased salt concentrations lead to changes in the cell wall composition, which, in turn, activate the salt stress response (Gigli-Bisceglia et al., 2022). Since the salt tolerance of the *PpFLOT*-OEX lines is severely impaired, the relative conductivity for all *PpFLOT*-OEX lines and a WT control was measured during cold stress. Observations of the relative conductivity in a temperature range of 0 °C to −7 °C for all tested genotypes revealed that the electrolyte leakage, and in turn, relative conductivity, increased in correlation with increased *PpFLOT* expression (Figure 11A). However, the variances between the measured replicates were too high to make a statistically significant statement on the data distribution. LC-MS analysis of the lipid profile of Δ*PpFLOT-1* and all *PpFLOT*-OEX lines revealed significant changes in their respective lipid levels with 27 differentially accumulated lipid classes in Δ*PpFLOT-1*, 13 in *PpFLOT*-OEX1, 17 in *PpFLOT*-OEX2 and 26 in *PpFLOT*-OEX3 (Supplementary Table 6). Interestingly, there was little overlap of the differentially abundant lipid classes among the four *PpFLOT* mutant lines. Only one lipid class, FA 20:4 (arachidonic acid), displayed significantly altered accumulation in all tested lines (Figure 11B, Supplementary Table 6). When comparing the *PpFLOT*-OEX lines, only digalactosyldiglyceride (DGDG) 34:6 and arachidonic acid were found to be differentially abundant in all lines (Figure 11B, 11C, Supplementary Table 6), with DGDG 34:6 decreasing and arachidonic acid increasing in the three overexpression lines correlating with the level of *PpFLOT* expression (Figure 11C). Similar to the results of the pigment and protein profile analyses, *PpFLOT*-OEX3 showed the greatest alterations in its lipid profile with 10 additional differentially accumulated lipid classes compared to the other *PpFLOT* mutant lines. Interestingly, when we looked for lipid classes that display altered levels in at least one *PpFLOT*-OEX line and show a change in the abundance correlating with the *PpFLOT* expression, we found one linolenic acid (FA 18:2) derivate displaying an increase correlated with the amount of PpFLOT. Similarly, the FA heptadecane acid (FA 17:0), ethyl linoleate (FA 20:2), heneicosanoic acid (FA 21:0), and triacontanoic acid (FA 30:0) showed their lowest abundance in the *PpFLOT*-OEX1 and their highest abundance in *PpFLOT*-OEX3 (Figure 11C). In addition to the FA, the glycerophosphoglycerol (PG) PG 34:2, glycerophosphocholines (PC), and phosphatidylethanolamine (PE) displayed similar behavior with the exception of PC 34:6 and PE 36:2, whose levels decreased with increasing *PpFLOT* expression (Figure 11C). Besides decreased DGDG 36:4 levels, MGDG 34:6 and 36:4 also displayed a low abundance. However, MGDG 36:4 shows higher levels in *PpFLOT*-OEX3 compared to *PpFLOT*-OEX1 and 2. This, combined with the general increase in the arachidonic acid pool, demonstrates the strongest effect of increasing *PpFLOT* expression on the lipid profile. These changes in the lipid profiles in the *PpFLOT*-OEX lines indicate alterations in composition of thylakoid membranes, attributed to variations in the abundance of DGDG and MGDG classes, which are primary constituents of thylakoid membranes. Additionally, the levels of FAs, crucial for cell wall component biosynthesis, potentially impacting the cell wall composition and subsequently, the sensitivity of these lines to abiotic stress, appear to be influenced by *PpFLOT* expression. Furthermore, the accumulation of arachidonic acid in *PpFLOT*-OEX plants suggests a potential role in plant defense mechanisms, as low concentrations can induce systemic resistance against pathogens, while high concentrations may lead to necrosis and phytoalexin accumulation (Dedyukhina et al., 2014). Arachidonic acid serves as precursor for oxylipins, oxygenated derivatives involved in plant defense, implying potential alterations in oxylipin biosynthesis and pathogen response, thus highlighting the involvement of PpFLOT not only in abiotic stress responses but also in pathogen defense mechanisms (Blée, 2002).

**Figure 11:**
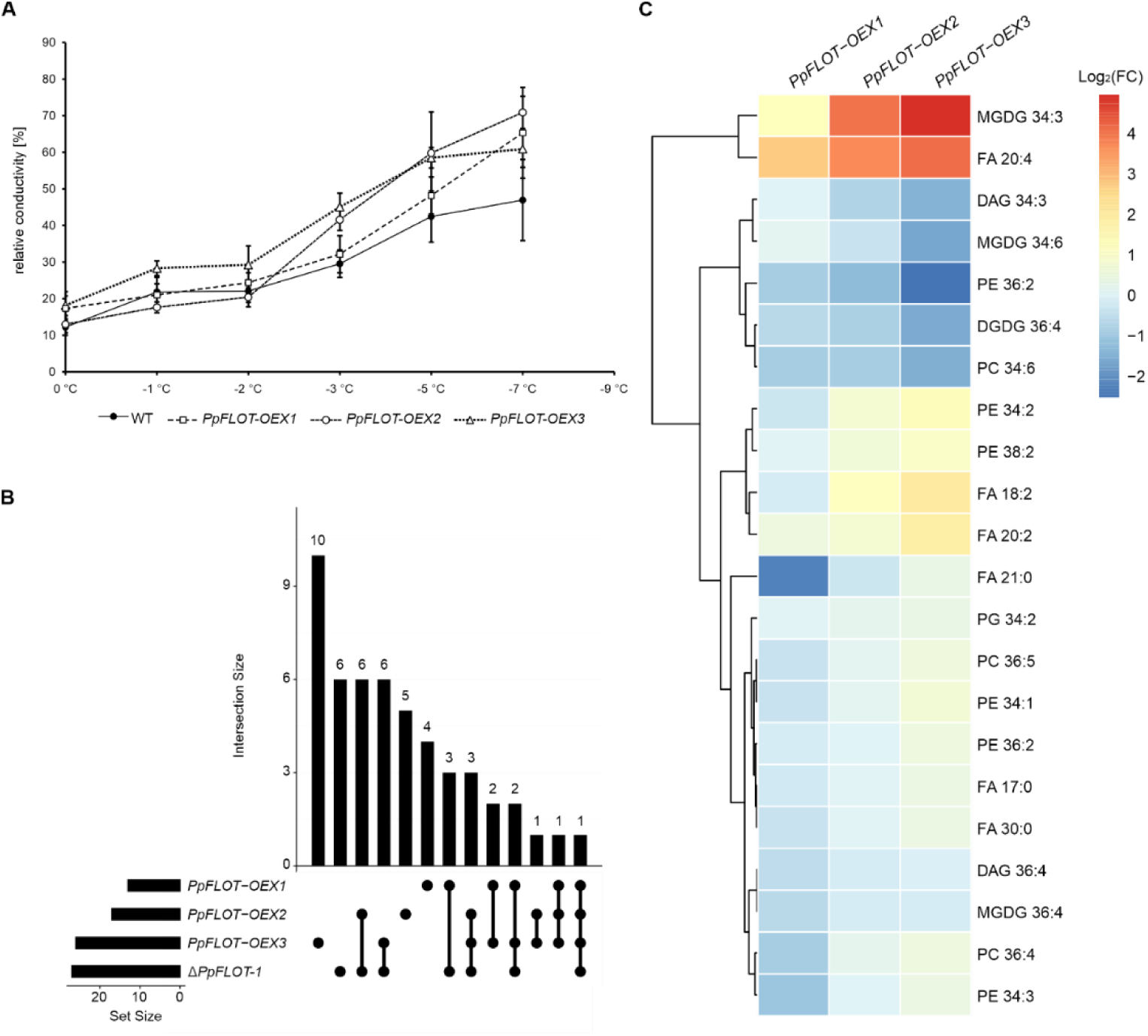
Analysis of changes in the lipid profile of *PpFLOT* mutant lines. (A) Conductivity measurements of 5 mg WT and *PpFLOT*-OEX gametophores cultivated for 11 weeks on solid medium at decreasing temperatures. Mean values ± SEM (n =3) of the relative conductivity are shown. (B) UpSet plot depicting the overlap of all identified differentially expressed (DE) lipids between Δ*PpFLOT-1* and all *PpFLOT*-OEX lines compared to the WT. Lipid levels are labeled as differentially accumulated when Student’s *t*-test using WT expression as control yielded p < 0.05 and the permutation-based false discovery rate (FDR) q < 0.05. (C) Clustered heatmap showing the log_2_(FC) of DE lipids of all *PpFLOT*-OEX lines. Lipids that show altered abundance in at least one line correlating to the amount of PpFLOT are displayed.

### *PpFLOT*-OEX affects grana stack assembly and enlarges the thylakoid lumen

Since PpFLOT localizes to the thylakoid membrane and alterations in thylakoid-membrane related proteins were observed in response to changes in the *PpFLOT* expression, we hypothesized that chloroplast structure might be impacted by *PpFLOT* overexpression. Transmission electron microscopy (TEM) analysis of chloroplast structures revealed significant deviations in the *PpFLOT*-OEX lines compared to WT and Δ*PpFLOT-1*, characterized by disordered thylakoid membranes, disrupted grana stacks, and enlarged thylakoid lumens (Figure 12). Interestingly, similar thylakoid alterations have been reported under high-light stress in Chlorophyta species and in seed plants during cold stress adaptation and salt treatment (Gorelova et al., 2019; Venzhik et al., 2019). Furthermore, examination of cell wall structure revealed a potential increase in cuticle layer density with increasing PpFLOT levels (Figure 12), possibly indicating changes induced by *PpFLOT* expression, supported by observed alteration in relative conductivity and fatty acid abundance in *PpFLOT*-OEX lines.

**Figure 12:**
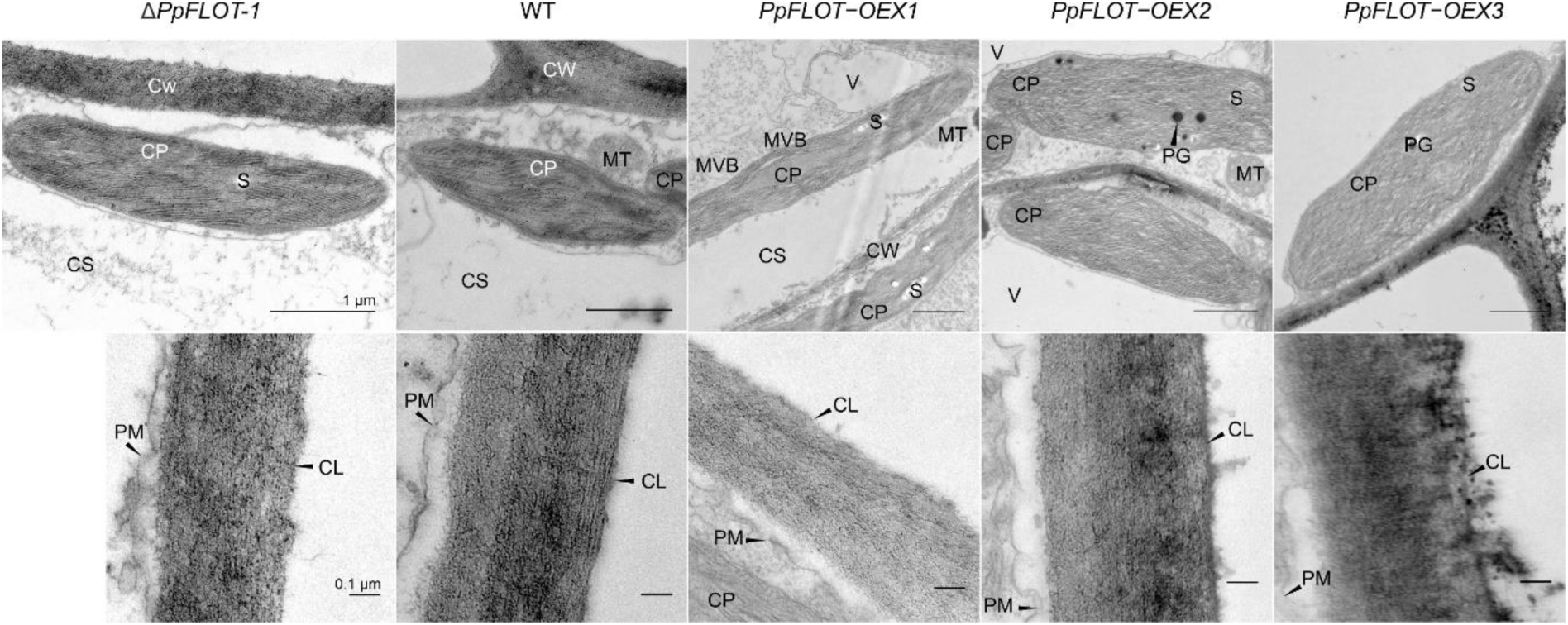
Altered *PpFLOT* expression leads to changes in the chloroplast thylakoid structure. Transmission electron microscopy (TEM) images of complete chloroplasts (upper panel) in protonema cells of Δ*PpFLOT-1*, WT and all three *PpFLOT*-OEX lines, along with close-up images of the cell wall in the respective lines (lower panel). Structural abbreviations include: chloroplast (CP), cytosol (CS), mitochondria (MT), cell wall (CW), plasma membrane (PM), vacuole (V), starch granula (S), plastoglobuli (PG), cuticle layer (CL) and multi vesicular bodies (MVB). Scale bars represent 1 µm (upper panel) and 0.1 µm (lower panel), respectively.

## Discussion

As mosses were among the first plants to adapt to life on land, the segregation of bryophyte FLOT in terms of function and localization from other plant FLOT is intriguing in the context of adapting to the new environment. Our observations suggest that both loss and overexpression of *PpFLOT* are less detrimental in the leafy gametophore than in the protonema life stage of *P. patens*, indicating the involvement of *PpFLOT* in the water-to-land transition. Our findings demonstrate that PpFLOT localizes to the plasma membrane in tobacco leaves, with no detectable presence in chloroplasts. This contrasts with its localization in *P. patens* and suggests an evolutionary divergence in FLOT targeting between bryophytes and seed plants. The absence of PpFLOT in tobacco chloroplasts raises questions about the molecular mechanisms governing its subcellular distribution, potentially influenced by species-specific regulatory elements such as post-translational modifications or protein-protein interactions. Whether at least one FLOT variant is positioned in the chloroplasts of other bryophyte, fern, or green algae species and what might have caused a shift in localization remains to be seen. To determine the most probable position of FLOT in the last common ancestor of land plants, further investigations in streptophyte algae, bryophytes, and seed plants are required. However, it is suggested that FLOT was acquired through horizontal gene transfer from fungi into ancient plant lineages (Ma et al., 2022), supported by the localization of the fungus FLOT homolog, FloA of *Aspergillus nidulans*, to plasma membranes (Takeshita et al., 2012). Thus, it is highly likely that a change in FLOT localization was driven by exaptation of FLOT function in *P. patens* and potentially in other bryophytes. At the protonema stage, we detected altered pigmentation in *PpFLOT*-OEX lines when grown in liquid culture under normal growth conditions. No such changes were detected in the leafy gametophores grown on solid medium, implying greater importance of FLOT under either submerged conditions or at the developmental stage of protonema cells. Indeed, changes in the cell shape were observed in response to *PpFLOT* overexpression, reminiscent of ABA-induced brachycytes that develop in *P. patens* as a survival mechanism in response to abiotic stress conditions (Arif et al., 2019). However, no significant changes were detected in the transcript expression of the ABA biosynthesis rate-limiting enzyme *PpNCED* (Hauser et al., 2011).

The accumulation of ABA levels to counter the increased *PpFLOT* levels is unlikely to be the only source of these changes, especially since 100 µm ABA is necessary to induce brachycytes (Arif et al., 2019). However, accumulation of PpLHCB2 correlated with increased PpFLOT levels. A study in 2012 showed that altered *LHCB* expression can influence ABA sensing and signaling (Xu et al., 2012), potentially affecting the ABA response in *PpFLOT*-OEX lines without impacting the ABA biosynthesis rate. Meanwhile, in *M. truncatula*, increased expression of *MtFLOT2* and *4* is crucial during nodule initiation upon infection with nitrogen-fixing bacteria (Haney and Long, 2010), and artificially activating CCaMK-IPD3 module in *P. patens* results in constitutively developed brood cells (Kleist et al., 2022). The CCaMK-IPD3 module is activated by symbiotic infection and oscillating calcium signals (Lévy et al., 2004; Miller et al., 2013; Kleist et al., 2022), suggesting that the formation of round, brachycyte-like cells in response to *PpFLOT* overexpression may involve slightly increased ABA levels and changes in Ca^2+^ signaling. Additionally, slight changes in *PpCRY1b* expression were detected, and since Ca^2+^ waves in *P. patens* can be induced by light and altered by changes in cryptochrome expression (Tucker et al., 2005), the circadian-regulated putative scaffolding protein PpFLOT might be involved in light-dependent Ca^2+^ signaling. External signals such as light-dark cycle, salt, mannitol treatment, and oxidative stress can influence Ca^2+^ dynamics between plastids and the cytosol (Sello et al., 2016; Martí Ruiz et al., 2020; Navazio et al., 2020).

It has been proposed that FLOT regulates the positioning of *A. thaliana* AQUAPORINE PIP1;2 (AtAPIP1;2) by initiating lipid rafts since AtAPIP1;2 colocalizes with AtFLOT1 (Browman et al., 2007; Li et al., 2011; Martiniere and Zelazny, 2021). Thus, by altering the composition of the thylakoid membrane, FLOT may putatively regulate the positioning of ion channels. For instance, PpFLOT might be involved in altering membrane fluidity or colocalize with Ca^2+^ channels, determining their position in thylakoid membranes. Moreover, Ca^2+^ transporters might not be the only ion channels affected by this. Our transcript expression analysis of the copper transport and HMA protein *Pp3c3_6890* showed a different response to ABA and salt treatment in *PpFLOT*-OEX1 than in both the WT and Δ*PpFLOT-1*. The GO-term analysis of all identified significant protein groups between WT and all *PpFLOT*-OEX lines also detected an effect of PpFLOT abundance on copper ion binding molecular function. Maintaining Cu homeostasis in chloroplasts is crucial, as Cu is a cofactor for the electron transporter plastocyanin (PCY), polyphenol oxidases (PPO), and Cu/Zinc superoxide dismutase (Cu/ZnSOD). PPO and Cu/ZnSOD participate in biotic and oxidative stress protection, respectively (Aguirre and Pilon, 2015; Printz et al., 2016; Schmidt et al., 2020). Interestingly, no significant change in the abundance of these Cu-dependent proteins was detected. Nonetheless, the expression levels of other copper transporters similar to *Pp3c3_6890*, might be affected, leading to changes in the copper homeostasis in *PpFLOT*-OEX lines. Further examination is necessary to determine if PpFLOT influences both Ca^2+^ signaling and copper transport.

As a scaffolding protein, PpFLOT is likely involved in multiple pathways and can oligomerize with multiple proteins (Garbett and Bretscher, 2014; Daněk et al., 2016). However, overexpression of *PpFLOT* impairs the high tolerance of *P. patens* against salinity and osmolarity stress (Frank et al., 2005), suggesting an additional function as a negative regulator during salt stress response. In response to long-term salt treatment, *PpFLOT* expression in WT is suppressed, likely due to elevated ABA levels, since ABA is a signaling molecule initiating the abiotic stress response (Hauser et al., 2011) and suppressing *PpFLOT* expression. Interestingly, PpTRX-Y2 and PpHSP70-1 are both suppressed in correlation to elevated PpFLOT levels. AtHSP70-1 participates in abiotic stress response regulation in *A. thaliana* (Cazalé et al., 2009; Leng et al., 2017), while AtTRX-Y2 is a known regulator of the light-dependent (Valerio et al., 2011; Seung et al., 2013; Geigenberger et al., 2017) and osmotic stress-induced starch degradation via amylases (Valerio et al., 2011). AtTRX-Y2 is also a crucial component of the antioxidative defense system in chloroplasts (Geigenberger et al., 2017). Hence, suppression of both proteins likely contributes to the observed altered salt tolerance. Although causality between PpFLOT expression and subsequent suppression of these two proteins is currently unknown, we can speculate about their relationship to PpFLOT. For example, HSP-70 proteins in plants drive protein translocation into organelles, including mitochondria and plastids (Shi and Theg, 2010; Berka et al., 2022). Downregulation of PpHSP70-1 might be a way for the cells to impede PpFLOT transportation in the chloroplasts of *PpFLOT*-OEX lines. Furthermore, AtTRX-Y2 reduces antioxidant enzymes, including peroxiredoxins (Collin et al., 2004; Jurado-Flores et al., 2020), glutathione peroxidases, and methionine sulfoxide reductases (Vieira Dos Santos et al., 2007; Laugier et al., 2013; Vanacker et al., 2018). Thus, changes in PpTRX-Y2 levels likely lead to the elevated ROS levels in response to elevated PpFLOT protein expression. Previous studies also showed that the LON DOMAIN-CONTAINING PROTEIN 1 suppresses TRX-Y2 activity and regulates ROS levels by controlling TRX-Y2 activity in *A. thaliana* (Shin et al., 2020). PpFLOT might putatively activate a similar protein in *P. patens,* and thus its overexpression leads to suppression of PpTRX-Y2 and increased ROS levels.

Upon detection of pathogens, ROS production within the chloroplast increases that activates stress signaling pathways to induce the plant defense against pathogens (Bleau and Spoel, 2021). Overexpression of PpFLOT leads to the accumulation of ROS, which is linked to the pathogen defense response (Hernández et al., 2016; Bleau and Spoel, 2021). Therefore, PpFLOT may have a putative role in the pathogen defense. In *A. thaliana*, treatment with flg22 alters the mobility of AtFLOT1, and AtFLOT1 overexpression increases callose deposition. Flg22 also induces AtFLOT aggregation (Yu et al., 2017; Junková et al., 2018). Our findings suggest that the cuticle layer and cell wall composition in the *PpFLOT*-OEX lines are altered due to an altered fatty acid profile. These changes of the cell wall components may be due to increased PpEXPA9 and PpPME53 accumulation in response to PpFLOT activated or guided pathogen defense mechanisms in *P. patens*. For example, an increase in Ca^2+^-dependent PME activity can lead to cell wall remodeling during abiotic stress response, and pectin fragments can be used as damage-associated signals (Shin et al., 2021). Moreover, PME activity increases after pathogen treatment in *A. thaliana* (Bethke et al., 2014). Increased PpFLOT expression not only leads to changes in the cell wall composition, but also to a higher abundance of linoleic acid derivatives and the accumulation of arachidonic acid. Pathogen defense signaling pathways depend on unsaturated FA (UFA), such as linoleic acid derivates (He and Ding, 2020). These UFA function not only as constituents for components of the cuticle layer, such as cutin and suberin, but also as intermediates in the biosynthesis of jasmonates and other active biomolecules of pathogen defense (Resemann et al., 2019; He and Ding, 2020). During pathogen defense, one way to counter the rising ROS is the oxidation of C18 UFAs into oxylipins, which themselves are building blocks of jasmonate biosynthesis (Blée, 2002; Resemann et al., 2019; He and Ding, 2020). The increase in the arachidonic acid pool further supports this hypothesis. Arachidonic acid is an elicitor of plant pathogen defense and depending on its abundance, its accumulation can also lead to either systemic resistance or to accumulation of phytoalexins and necrosis of plant tissues (Dedyukhina et al., 2014). There is a distinct possibility that the detected changes in coloration of *PpFLOT*-OEX cultures are the result of increased necrotic events. Interestingly, we also detected increased levels of PpSEN1 in all three *PpFLOT*-OEX lines. SEN1 not only shows increased expression in *A. thaliana* in response to infection events but also triggers the senescence response resulting in necrosis upon pathogen infection (Schenk et al., 2005; Fernandez-Calvino et al., 2016). A senescent phenotype in *P. patens* can also lead to changes in the FA composition, including arachidonic acid (Chen et al., 2020). Changes in culture coloration of *PpFLOT*-OEX lines might thus be attributed to necrosis rather than changes in pigments of the xanthophyll cycle. Even though these pigments displayed an overall low abundance in *PpFLOT*-OEX lines, changes in coloration due to altered expression of the unidentified pigments are still a distinct possibility. Curiously, no changes in coloration were detected in the gametophores of *PpFLOT*-OEX lines.

We suggest that the increased production of ROS triggers SEN1 accumulation (Schenk et al., 2005) and may cause enhanced hypoxia in chloroplasts (Pucciariello and Perata, 2021), which cannot be compensated by *P. patens* protonema cells in an already anoxic environment. The molecular changes that enable 3D growth of *P. patens* may help the plant to counteract this effect. For example, *PpFLOT* expression in the WT gametophore is lower compared to its expression in rhizoids, caulonema cells and protoplasts (eFP browser, https://bar.utoronto.ca/efp_physcomitrella/cgi-bin/efpWeb.cgi) (Winter et al., 2007).

Other regulators of systemic acquired resistance (SAR) in response to pathogen infection include DGDG and MGDG (Gao et al., 2014). DGDG promotes SAR by regulating NO and salicylic acid synthesis, while MGDG regulates signals downstream of NO-guided SAR, such as azelaic acid and glycerol-3-phosphate (Gao et al., 2014). Although proteomics and metabolomics analysis did not detect any DE of these signals, overexpression of *PpFLOT* severely altered MGDG and DGDG levels. Therefore, overexpressed MGDG species may be beneficial during the pathogen response of *PpFLOT*-OEX lines.

In addition to its proposed contribution to pathogen defense, PpFLOT seems to be involved in chlorophyll biosynthesis. An increase in PpLHCB2 protein levels was detected in correlation with PpFLOT levels, and the two *PpFLOT*-OEX lines with the highest PpFLOT and PpLHCB2 expression also showed elevated chlorophyll levels. PpLHCB2 could potentially be a direct interaction partner of PpFLOT, but it is also a chlorophyll-binding molecule, and its overexpression could be due to a need for binding excessive chlorophyll (Eberhard et al., 2008; Johnson, 2016). Even slight changes in the expression of one LHCB influence the expression and overall composition of the light-harvesting antenna of PSII (Xu et al., 2012). Furthermore, LHCB expression takes part in regulating both ABA and ROS homeostasis (Xu et al., 2012), and high ABA levels can induce LHCB expression (Liu et al., 2013). Interestingly, ROS is one of the signaling molecules used for plastid to nucleus communication the retrograde signaling (Eberhard et al., 2008; Li and Kim, 2022). *GLK2*, a gene involved in retrograde signaling and chlorophyll biosynthesis regulation (Yasumura et al., 2005; Kim et al., 2023; Lee et al., 2023), is upregulated in response to *PpFLOT* knockout, but downregulated in *PpFLOT*-OEX lines. This gene acts as a transcription factor for photosynthetic genes, and its knockout in *A. thaliana* leads to diminished chlorophyll content, while *AtGLK2* overexpression results in higher chlorophyll levels (Yasumura et al., 2005; Kim et al., 2023). Both knockout and overexpression of *PpFLOT* lead to changes in the expression of *PpGLK2*, suggesting a role of PpFLOT in chlorophyll biosynthesis. However, no changes in the protein levels of known factors participating in chlorophyll biosynthesis could be detected, suggesting a regulating function rather than a direct involvement. Since the chlorophyll content is decreased in Δ*PpFLOT-1* and increases with the *PpFLOT* expression, the anticorrelating expression of *PpGLK2* to the respective chlorophyll content suggests an attempt to regulate PpFLOT-dependent changes in the chlorophyll biosynthesis via *PpGLK2*. We propose that PpFLOT most likely participates in the recruitment of protein complexes involved in chlorophyll biosynthesis to the thylakoid membrane. Due to the circadian nature of PpFLOT and its inducibility by darkness, one of its functions might contribute to the light-dependent regulation of chlorophyll biosynthesis. For instance, FLUORESCENT IN BLUE LIGHT (FLU) suppresses the chlorophyll biosynthesis in the dark by inactivating the GLUTAMYL-tRNA-REDUCTASE (GluTR) and repressing the 5-aminolevulinic acid (ALA) synthesis (Meskauskiene and Apel, 2002; Kauss et al., 2012; Fang et al., 2016; Hou et al., 2019; Wittmann et al., 2021). ALA is an intermediate compound during chlorophyll biosynthesis, and its decline during the night prevents the accumulation of phototoxic products (Wittmann et al., 2021). How GluTR is recruited into a complex together with the membrane-bound FLU and other components of the Mg^2+^ branch of the chlorophyll biosynthesis is unclear (Wittmann et al., 2021). Our findings suggest that at least in *P. patens*, PpFLOT might fulfill such a role. Since, in this case, the activity of FLU would be dependent on PpFLOT abundance, chlorophyll biosynthesis might be suppressed in the *PpFLOT*-OEX1, leading to the detected lower chlorophyll levels. In this case, both *PpFLOT*-OEX2 and 3, as well as the ΔPpFLOT-1 lines, might depend on other ways to regulate the chlorophyll synthesis under changing light conditions, leading to higher and lower chlorophyll levels, respectively. However, concrete evidence supporting this hypothesis is still lacking. It is possible that PpFLOT influences chlorophyll biosynthesis in another way. However, altered chlorophyll levels are the most likely explanation for the detected changes in the Y(II) of *PpFLOT*-OEX1 and 2 and the NPQ/4 of Δ*PpFLOT-1*. In Δ*PpFLOT-1*, excess energy that cannot be processed due to inefficient chlorophyll levels is redistributed and redirected into non-photochemical quenching (Eberhard et al., 2008). On the other hand, an excess of chlorophyll leads to a higher photorespiration in *PpFLOT*-OEX lines. This increase in photorespiration most likely also leads to an increase in respiration, supported by the measured increase in PpNDUFS8 and PpNDUFA2/B8 expression. Previous studies showed that the thylakoid lumen enlarges in response to abiotic stresses (Gorelova et al., 2019; Venzhik et al., 2019) to increase the travel distance and diffusion rate of electron carriers to the photosystems, thus slowing down the photosynthesis rate (Mullineaux, 2008; Kirchhoff et al., 2011; Jarvi et al., 2013; Gorelova et al., 2019). Such a reaction would also explain the observed lumen enlargement in *PpFLOT*-OEX chloroplasts. Increased PpFLOT expression most likely contributes to this effect.

Detected changes in the lipid composition of thylakoid membranes could also explain the increase in thylakoid lumen and structural changes in these membranes. Thylakoid membranes mainly consist of the glycolipids MGDG, DGDG, and sulfoquinosyl-diacylglycerol (SQDG), with MGDG making up more than 50 % of the total lipid content, while DGDG makes up about 30 % (Rast et al., 2015; Garab et al., 2016). Unlike DGDG, MGDG is a non-bilayer lipid that is forced into a bilayer by interaction with LHCII proteins (Garab et al., 2016). PpFLOT may be involved in the localization or transport of PpLHCB2 to the PS II antenna complexes along the thylakoid membranes, which could affect the structure of the thylakoid membrane. Meanwhile, DGDG is the main driver behind the bilayer formation, and changes in the MGDG/DGDG ratio affect both membrane organization and protein complex stability (Pribil et al., 2014; Rast et al., 2015; Garab et al., 2016). Although the exact mechanism behind increased PpFLOT levels and subsequent changes in thylakoid membrane composition are unknown, it can be assumed that this change in lipid composition contributes to the detected alterations in the thylakoid structure of the *PpFLOT*-OEX lines. Interestingly, increased levels of MGDG can also lead to a higher photosynthetic activity (Zhou et al., 2009; Pribil et al., 2014). One way that PpFLOT could adjust the lipid composition of thylakoid membranes directly is by participating in the formation of stromal vesicles that transport lipids generated in the chloroplast envelope to the thylakoid membranes.

Vesicle formation has been observed in proplastids and developing chloroplasts (Pribil et al., 2014; Mechela et al., 2019), potentially playing a crucial role in early thylakoid membrane development (Mechela et al., 2019). While AtFLOT1 has been implicated in clathrin-independent endocytosis (Li et al., 2012; Cao et al., 2020) and vesicle formation, it is challenging to extrapolate the molecular function of PpFLOT from AtFLOT due to their differing locations. Besides, the absence of evidence for chloroplast-related FLOT in other plant species, coupled with the lack of observable changes in thylakoid structure in Δ*PpFLOT-*1, suggests that FLOT may not significantly contribute to chloroplastic vesicle formation outside of *P. patens*.

It is difficult to distinguish between direct and indirect effects of PpFLOT overexpression since no claims can be made about its direct interaction partners. However, this study has revealed a few important points. PpFLOT is more versatile than previously assumed, and while its association to thylakoid membranes may be unique to bryophytes, the differences in the gametophore and protonema phenotype suggest a change in function during the transition of plant terrestrialization. Although the exact function of PpFLOT is still a mystery, this study has shed light on a few pathways in chloroplast metabolism where PpFLOT appears to be crucial. High concentrations of PpFLOT are detrimental during salt adaptations and to cope with elevated osmotic pressure. However, evidence gathered suggests a potential role of PpFLOT in pathogen defense response, chlorophyll biosynthesis, and involvement in Ca^2+^ signaling (Figure 13).

**Figure 13:**
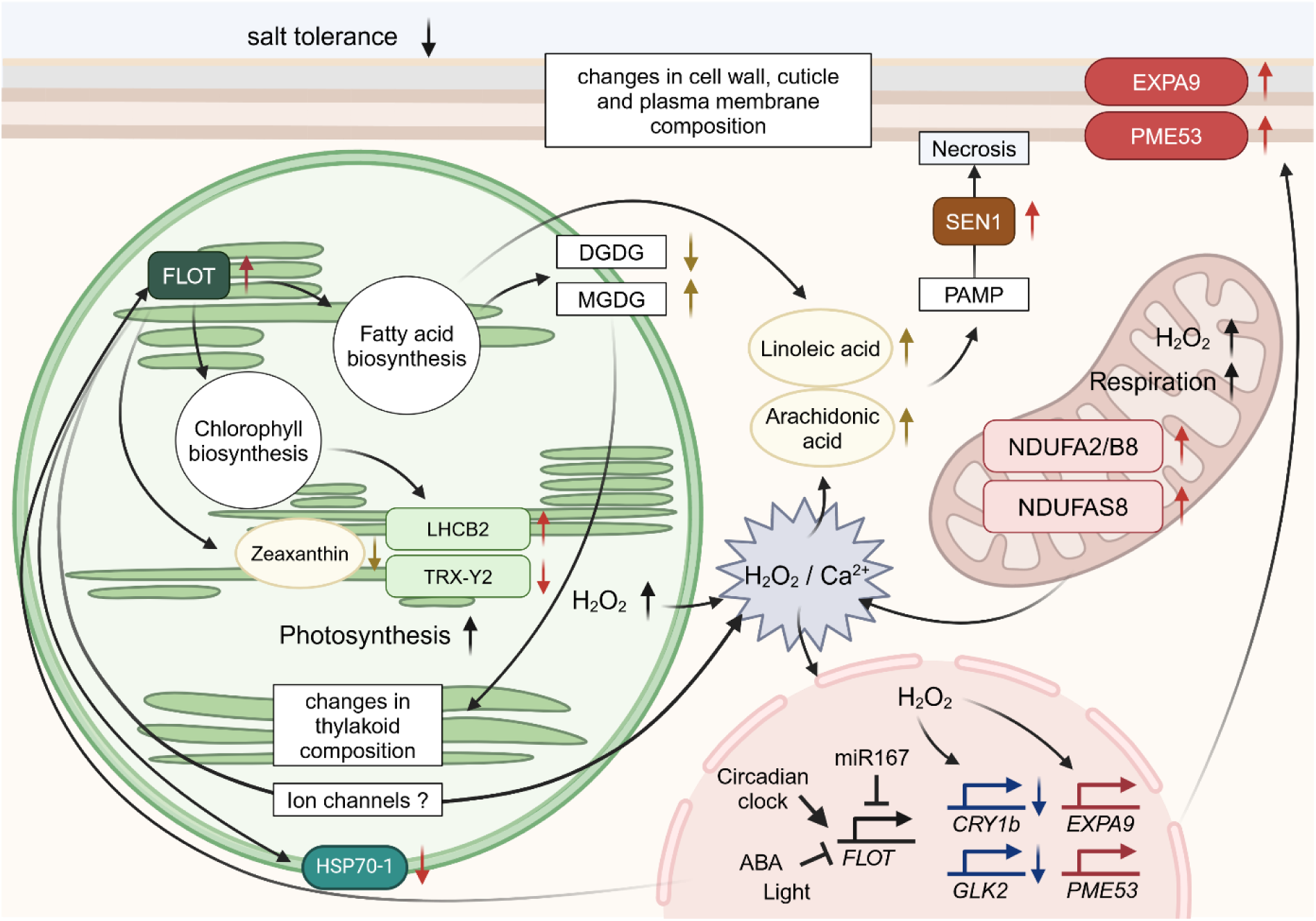
Effect of *PpFLOT* overexpression in protonema cells. The expression of *PpFLOT* is regulated by the circadian clock, ABA, light stimuli, and potentially by miR167. Elevated PpFLOT levels in chloroplasts alter fatty acid biosynthesis, increase chlorophyll biosynthesis, decrease PpHSP70-1 levels, and likely impact ion channel activity or localization. Moreover, high PpFLOT levels increase PpLHCB2 while reducing zeaxanthin and PpTRX-Y2 levels. Changes in fatty acid biosynthesis affect thylakoid membrane composition due to an increase in MGDG and a decrease in DGDG levels. These changes enlarge the lumen and impair grana formation in the thylakoid membrane, enhance photosynthesis, H2O2 production, and possibly Ca^2+^ signaling. Elevated photosynthesis affects metabolic processes including respiration, as evidenced by upregulated PpNDUFA2/B8 and NDUFAS8. H_2_O_2_ accumulation triggers retrograde signaling increasing the expression of *PpEXPA9* and *PpPME53* and suppressing the expression of *PpCRY1b* and *PpGLK2*. Increased PpEXPA1 and PpPME53 levels alter cell wall, plasma membrane, and cuticle composition. Together with low levels of PpHSP70-1, PpTRX-Y2, and the overall increased metabolic activity changes in these components are reducing the salt tolerance. Altered fatty acid biosynthesis leads to linoleic acid and arachidonic acid accumulation, potentially contributing to pathogen-associated molecular patterns (PAMP), including the accumulation of PpSEN1, leading to necrotic events. The arrows in the graph denote experiments validating changes in expression levels or protein abundance. Blue: qRT-PCR measurements, red: Proteomics analysis, yellow: Metabolomics analysis, black: PAM and DAB staining. This graphical representation was generated using BioRender.

## Material and Methods

### *PpFLOT::citrine* localization in *P. patens* protoplasts

The *PpFLOT* transcript (Pp3c3_21910) was tagged with citrine by cloning its CDS into a vector containing the Actin 5 promotor (ACT5_P), citrine sequence with required linkers for the tag (citrine) (Tian et al., 2004; Top et al., 2021), and a *nopaline synthase* terminator (NOS_T). The complete *PpFLOT* CDS was amplified using the oligonucleotides 5’-AGCTCTCGAGATGGCGTTCCATACCGC-3’ and 5’-TCTAGATCTGGCTTGGGGAAGCTTGG-3’, which generated a *PpFLOT* sequence flanked by two restriction enzyme sites for *Bgl*II and *Xho*I. This sequence was then ligated into a linearized Act5-citrineL vector using T4 ligase (Invitrogen) following manufacturer’s instructions. The generated *PpFLOT::citrine* construct was transformed into *P. patens* protoplasts to generate lines that transiently express *PpFLOT::citrine*. Images of *PpFLOT::citrine* protoplasts were captured three days after transformation with a Stellaris 5 Point Scanning Confocal Microscope (Leica Microsystems) equipped with a turnable white light laser (485 - 685 nm) and five power hybrid HyD S detectors. The imaging was performed using the immersion oil objective HC PL APO CS2 63x/1.40. Chlorophyll autofluorescence (chl) served as a localization marker, excited at 405 nm and recorded at 623 nm–750 nm while the fluorescent citrine protein (citrine) was excited at 512 nm and recorded at 524 nm–560 nm. Image processing was conducted using the LAS X Office software.

### PpFLOT::venus localization in *N. benthamiana*

The *PpFLOT* CDS was cloned into the pHKL0786 vector using Gibson Assembly (NEB), transformed into *Escherichia coli*, and confirmed by sequencing. For *Agrobacterium tumefaciens*-mediated expression, *Agrobacterium tumefaciens* GV3101 containing the pSOUP helper plasmid (Hellens et al., 2000) was co-injected with the 19k vector (Voinnet et al., 2003) and infiltrated as described by Waadt et al. (2014) with minor modifications. The HygR selection marker was replaced with a fast-red selection cassette, which expresses RFP under the control of the Olesin promoter in the seed coat. *A. tumefaciens* cultures transformed with pHKL0786-*PpFLOT::venus* and pSoup were electroporated at 1.66–1.9 kV, recovered in LB medium, and plated on selective LB agar for incubation at 28°C overnight. Transformed *A. tumefaciens* cultures were then used for transient transformation of *N. benthamiana* leaves.

Three days post-transformation, leaf samples were collected for fluorescence imaging. Confocal imaging of leaf tissue was performed using a Stellaris 8 Point Scanning Confocal Microscope equipped with a tunable white light laser (485–685 nm) and an HC PL FLUOTAR L 20x/0.40 DRY objective. Additionally, imaging was also conducted using a Stellaris 5 Confocal Laser Scanning Microscope (Leica, Wetzlar), equipped with a 405 nm diode and a supercontinuum White Light Laser (WLL), with fluorescence detection using a Power Hybrid Detector HyD S. For visualization, the recombinant protein fused to mvenus (YFP) was excited at 514 nm, and emission was recorded between 520–580 nm. Chlorophyll autofluorescence was excited at 405 nm, with emission detected in the 623–813 nm range. In the Stellaris 8 system, chlorophyll autofluorescence was excited at 440 nm and recorded at 670–685 nm, while venus fluorescence was excited at 510 nm and recorded at 539–550 nm. Image processing was conducted using LAS X Office software.

### Phylogenetic analysis

To analyze the evolutionary relationship of PpFLOT to FLOT homologs in other species we performed query searches with the full-length amino acid sequence (aa) of PpFLOT in multiple plant species. Reciprocal searches with the BLASTp function of Phytozome v13 (phytozome-next.jgi.doe.gov), UniProt (re_2024_01; uniprot.org) and MarpolBase (marchantia.info) databases revealed PpFLOT homologs in *A. thaliana*, *A.us officinalis*, *A. filiculoides*, *C. purpureus*, *C. braunii*, *C. subellipsoidea*, *K. nitens*, *M. polymorpha*, *M. truncatula*, *O. sativa*, *S. cucullata*, *S. polyrhiza*, *Synechocystis* sp. PCC 6803, *Z. mays*, and *Z. marina*. The full-length aa sequences of the detected PpFLOT homologs were aligned using CLC Genomics Workbench v20.0.4 (Quiagen), followed by phylogenetic analysis employing the Maximum Likelihood method with 1000 bootstrap replicates and the Jones-Taylor-Thornton method as the substitution model. The phylogenetic tree was generated using the Nearest-Neighbor-Interchange method in MEGA X (Kumar et al., 2018).

### Generation of *PpFLOT* knockout and overexpression lines

To create Δ*PpFLOT* lines, exon 2 of the *PpFLOT* was replaced with a *neomycin phosphotransferase II* (*nptII*) selection cassette through homologous recombination. A *PpFLOT* CDS construct with the embedded *nptII* cassette was generated using Gibson assembly (NEB) (oligonucleotides listed in Supplementary Table 1) and then cloned into pJET1.2/blunt (Thermo Fisher Scientific) vector. After amplification, the construct was cut out of the plasmid via a *BglII* restriction site, separated from the pJET1.2 backbone by agarose gel electrophoresis and purified. The construct was transformed into *P. patens* protoplasts, and transformed plants were selected by growing them on medium supplemented with 50 µg/ml G418 sulfate. The correct integration of the *nptII* cassette was confirmed by amplifying the full-length *PpFLOT* genomic sequence containing the *nptII* cassette and the predicted integration sites by PCR (oligonucleotides listed in Supplementary Table 1). Loss of the *PpFLOT* transcript was confirmed by qRT-PCR using primers located within the *PpFLOT* CDS (oligonucleotides listed in Supplementary Table 1).

To create *PpFLOT* overexpression lines, the entire *PpFLOT* CDS was ligated into a vector carrying an ACT5_P sequence, a NOS_T sequence and a hygromycin selection cassette. ACT5_P-controlled *PpFLOT* CDS fragments encoding additional hygromycin resistance were transformed into *P. patens* protoplasts. The transformed plants were selected by cultivating them on hygromycin-containing medium and pre-screened by detecting the hygromycin resistance cassette by PCR (oligonucleotides listed in Supplementary Table 1) and by amplifying ACT5_P controlled *PpFLOT* CDS with primers spanning from the ACT5_P region to the NOS_T region of the newly inserted *PpFLOT* sequence. The overexpression of PpFLOT was confirmed by amplifying of the *PpFLOT* transcript from cDNA using Pp_Flot_fwd and Pp_Flot_rev primers (Supplementary Table 1). The strength of the overexpression was determined by qRT-PCR with the primers Flot_qRT_fwd and Flot_qRT_rev (Supplementary Table 1).

### Confirmation of single integration lines by Southern blot analysis

Genomic DNA was extracted following the CTAB DNA extraction protocol as described by Inglis et al. (2018). From each Δ*PpFLOT* line, 10 µg DNA was digested overnight with X*hoI* and *NcoI*. After digestion the genomic DNA was separated by agarose gel electrophoresis and then transferred to an Hydrobond-N+ membrane (GE Healthcare) using an adapted alkalic desaturating blotting following the capillary method (Brown, 2001). After washing and pre-hybridization of the membrane, the hybridization probe was radioactive labeled with the Prime-a-Gene® Labeling System (Promega). A complementary probe against the *nptII* selection cassette was generated by PCR with the primers 5’-TCCATCATGGCTGATGCAAT-3’ and 5’-GGCGATACCGTAAAGCACGA −3’ using the *nptII* containing pUC-*nptII* vector (Top et al., 2021) as template. After overnight hybridization and subsequent washing of the blot, radiation was detected with phosphor image screen for at least 4 h before scanning the screen with the Typhoon Trio Varible Mode Imager (Amersham Biosciences). The brightness of the image was adjusted with the open-source program ImageJ (Schneider et al., 2012).

### Plant material cultivation and phenotypic analysis

*Physcomitrium patens* ssp. *patens* (Hedwig) ecotype “Gransden” 2004 and all generated mutant lines were grown under long-day conditions (16 h light/ 8 h dark) at 23 °C and a light intensity of 85–100 µmol/m^2^s. Axenic cultivation was performed in liquid or on solid medium as previously described (Frank et al., 2005). Liquid cultures were homogenized and transferred to fresh media every 14 d. Plants were cultivated on solid medium supplemented with 250 mM NaCl, 300 mM NaCl, 700 mM mannitol, 10 µM 2-*cis*,4-*trans*-abscisic acid (ABA)(Sigma-Aldrich), 10 µM 1-naphthylacetic acid (NAA) (1mg/ml, Sigma-Aldrich), 10 µM 6-y-y-(dimethylallylamino)-purine (2-ip) (Duchefa), respectively. For the phenotypic analysis, liquid cultures of all lines were adjusted to an equal density of 100 mg dry weight/L, and 5 µl of the generated knockout, overexpression, and WT control lines were spotted on solid medium. The phenotype assay was performed in triplicates and culture growth was observed and documented for 8 weeks. Pictures were taken with a SMZ 1500 stereomicroscope and a DS-U3 camera (Nikon).

### Growth assay

To assess changes in the growth rate between the WT and the generated Δ*PpFLOT* lines under control conditions and salt treatment, protonema cultures were adjusted to an equal density of 100 mg dry weight/L in liquid medium, with and without supplementation of 250 mM NaCl. Cultures were monitored for 14 d, with dry weight measurements taken every two to three days. Culture pigmentation was documented using a DS-U3 camera (Nikon). These measurements were conducted in three independent experiments.

### Analysis of circadian expression control of *PpFLOT*

To investigate whether *PpFLOT* expression is influenced by light or daytime, WT subcultures of 30 mg dry weight protonema were cultivated for three days under different light conditions: long-day 16 h light/8 h dark photoperiod (LD 16h:8h), continuous light (CL), and complete darkness (D). After three days, protonema cultures were harvested at 4 h intervals over a 24 h period (0, 4, 8, 12, 16, 20 and 24 h). Light intensity and temperature conditions were kept to standard cultivation conditions. RNA extraction, sample preparation, and qRT-PCR were performed using EVAGREEN®DYE (Biotium) containing reaction mixes as previously described by Arif et al. (2022). To detect time-dependent changes in expression and differences between light treatments, normal distribution and equality of variances of the data sets were confirmed using Shapiro-Wilk and Levene tests, respectively, before ANOVA analysis. Statistically significant differences in *PpFLOT* expression among light treatments and time points were determined using Tukey’s HSD test. Cosinor analysis was additionally performed to determine potential rhythmicity in *PpFLOT* expression. All statistical analyses were performed in R (v. 4.2.2) (R Core Team, 2023) using the packaged “rstatix” (Kassambara, 2023b) and “cosinor2” (Mutak, 2018). Results were considered significant when p < 0.05.

### Gene expression analysis after ABA and salt treatments

Liquid cultures of WT, Δ*PpFLOT-1* and *PpFLOT*-OEX1 were adjusted to a density of 0.4 mg dry weight/ml. The cultures were then divided into three groups per line, untreated control cultures and cultures treated with 10 µM ABA and 250 mM NaCl, respectively. Samples were harvested from each line and treatment in triplicates at 0, 1, 2, 4, 8 and 24 h after treatment. RNA extraction and qRT-PCR were performed as described above and statistical ANOVA analysis was performed to determine statistically significant differences between the analyzed lines under control conditions and after the treatment. When p < 0.05 a Tukey’s HSD test was performed to identify the significantly different lines and the time points of treatment where these differences could be identified. The statistical analysis was conducted with R (v4.2.2) as previously described. MiRNA expression was analyzed using the stem loop PCR method according to Kramer (2011) following the protocol described by Tiwari et al. (2021) adapted for *P. patens* and using *PpEF1α* as cDNA synthesis control.

### Analysis of photosynthetic activity

Pulse-Amplitude-Modulation (PAM) fluorometry was used to measure the photosynthetic activity of WT, Δ*PpFLOT-1*, *PpFLOT*-OEX1, *PpFLOT*-OEX2, and *PpFLOT*-OEX3 during their gametophore and protonema life stages. For the analysis of photosynthetic activity in the gametophores, liquid cultures of all lines were standardized to a density of 100 mg/L and spotted in triplicate 5 µl spots on solid medium before cultivating them for three months to ensure densely grown moss colonies. Three independent experiments were prepared for the gametophores (n = 9 per analyzed line). To measure the fluorescence in protonema, liquid cultures of all lines were adjusted to a concentration of 1 mg dry weight/ml. Five ml of the liquid cultures were filtered through Miracloth (Merck) tissue and placed into six well plates (n = 9 per analyzed line). All plates were dark adapted for 3 h before PAM measurements were performed with the IMAGING PAM M-Series chlorophyll fluorescence imaging system (Walz GmbH) and the ImagingWinGigE V2.56zn software program. Actinic light of 450 nm (IMAG-Max/L) was used at intensity of 55 µmol/m²s photosynthetically active radiation (PAR) to mimic normal growth conditions and the saturation pulse was given for 240 ms to determine maximal fluorescence after dark-adaptation. After 40 s, the steady-state fluorescence (F) levels were determined. The steady-state F levels were measured at actinic light intensities of 55 µmol/m²s PAR in 20 intervals for 315 s. Since protonema cultures of *PpFLOT*-OEX3 displayed low fluorescence signals and fluorescence could not be detected with these parameters the measuring light intensity was adjusted, resulting in an actinic light intensity of 60 µmol/m²s PAR. Due to these adjustments, PAM measurements of *PpFLOT*-OEX3 protonema cultures were not considered during the statistical evaluation. PSII quantum yield [Y(II)] was calculated as (Fm’ -F)/Fm’ (Genty et al., 1989) and the non-photochemical quenching (NPQ/4) as ((Fm-Fm’)/Fm’)/4 (Kramer et al., 2004; Gao et al., 2022). To detect statistically significant differences between the analyzed lines, ANOVA analysis was performed, and when p < 0.05 than the statistical analysis was followed up with a Tukey’s HSD test to determine which lines show significant differences. Statistical analysis for all parameters was performed with R (v. 4.2.2) using the “rstatix” (Kassambara, 2023b) package, and graphs were generated with R using the “ggpubr” (Kassambara, 2023a) package.

### H_2_O_2_ detection by 3,3-Diamonobenzidine staining

Accumulation of H_2_O_2_ in all mutant lines was analyzed by harvesting 15 ml of WT, Δ*PpFLOT-1*, *PpFLOT*-OEX1, *PpFLOT*-OEX2, and *PpFLOT*-OEX3 protonema cultures grown under standard conditions. The harvested protonema was equally divided into samples treated with sterile MilliQ H_2_O (mock), 1 mg/ml 3,3-Diamonobenzidine (DAB) pH 3.8 (Rea et al., 2004), and medium (control) for each line before incubating the samples for 18 h at CL. Subsequently, the solutions of the mock and DAB samples were replaced with 99.9 % ethanol, boiled for 10 min, and stored in fresh ethanol, while the control remained untreated. Microscopy images of mock, DAB and untreated control of all lines were taken with an Axiophot light-microscope (Carl Zeiss Microscopy GmbH). Reddish-brown colorations are indicating the sites of H_2_O_2_ accumulation.

### Evaluation of chlorophyll content

To evaluate the chlorophyll content of WT, Δ*PpFLOT-1*, and all *PpFLOT* overexpression lines, 0.4 g protonema fresh-weight was harvested in triplicates for each line and immediately frozen in liquid nitrogen. The harvested plant material was homogenized with metal beads in a TissueLyser™ (Quiagen) to obtain a fine powder, and chlorophyll was extracted by adding 1.5 ml of 80% (v/v) acetone to all samples. The samples were then mixed at 300 rpm for 5 min before centrifugation at 14,000 *g* for 5 min to obtain cell free chlorophyll solution. Chlorophyll content was determined by measuring the absorbance of the supernatant at 645 and 663 nm. Subsequently, the supernatant was transferred back into the reaction tube containing the cell debris of the respective lines, and the samples were dried in a speed-vac centrifuge at room temperature overnight. Finally, the samples were dried for 2 h at 105°C to obtain the dry weight for all samples. The chlorophyll content was then calculated after the following method:

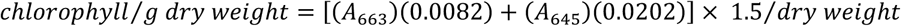

### Electrolyte leakage assay

To assess electrolyte leakage in WT and all *PpFLOT* overexpression lines, 5 mg of gametophores grown on solid medium for 8 weeks were collected without damaging the tissue and placed into clean round glass tubes. A clean, adjusted cell sieve was placed into each glass tube without damaging the harvested plant material to serve as physical barrier between the plant material and the electrode of the conductivity meter during measurements. Glass tubes were filled with 5 ml ddH_2_O and incubated in a cryostat for 1 h at 0 °C. Ice formation was induced by introducing a metal wire pre-cooled in liquid nitrogen to the water surface without disturbing the plant material. Samples were returned to the cryostat, and every 30 min, the temperature was decreased by 1 °C. Incubation was stopped for all lines in triplicates at 0, −1, −2, −3, −5, and −7 °C. Samples were left to thaw while gently shaking overnight at 4 °C until all ice crystals were completely dissolved. After adding 5 ml ddH_2_O samples were gently shaken additional 30 min before measuring their conductivity with an inoLab® Cond 7110 (Xylem Inc.) conductivity meter. Samples were autoclaved for 20 min, cooled for 30 min at 4 °C, and then allowed to adjust to room temperature by gently shaking at room temperature for 45 min before measuring total conductivity. The relative conductivity was calculated as follows:

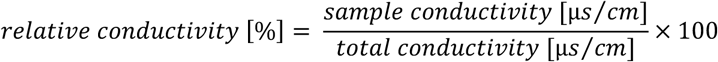

### Proteomics analysis

DE protein expression in *PpFLOT*-OEX1, *PpFLOT*-OEX2, and *PpFLOT*-OEX3 lines was analyzed by generating total protein extracts from 50 mg protonema (fresh weight) for these lines and the WT control in four replicates. Protein extraction and trypsin digestion were performed as described in (Marino et al., 2019), Liquid chromatography-tandem mass spectrometry (LC-MS/MS) was performed following the protocol described by Espinoza-Corral et al. (2023) with the exception that peptides were separated over a 90 min linear gradient of 5 – 80% (v/v) acetonitrile (ACN). Raw files were processed using the MaxQuant software version 2.1.0.0 (Cox and Mann, 2008), annotating detected peaks against the *P. patens* reference proteome (UniProt, www.Uniport.org) using the “match-between-runs” setting. Proteins were quantified via label-free quantification method (LFQ) previously described by Cox et al. (2014), and subsequent analysis utilized Perseus version 2.0.1.1. (Tyanova et al., 2016; Tyanova and Cox, 2018). Contaminants, reverse hits, and proteins identified solely by site modifications were excluded from further analysis. Only protein groups quantifiable by LFQ algorithm in at least three out of four technical replicates of one of the analyzed lines were used. LFQ intensities were log_2_-transformed, and missing values were imputed from a normal distribution using default settings. ANOVA tests in Perseus identified statistically significant protein groups between all tested lines (p < 0.05, permutation-based false discovery rate (FDR) q < 0.05). Z-scores of significant log_2_-transformed LFQ intensities were used to generate a clustered heatmap for easy comparison. GO-term enrichment analysis of differentially expressed protein groups (DEP) was performed using web based shinyGO v0.8 application (Ge et al., 2020). Additional Student’s *t*-tests between all *PpFLOT*-OEX lines and the WT control identified significantly differentially expressed protein groups (all p < 0.05, permutation-based FDR q < 0.05 and log_2_(LFQ intensities) < −1 and > + 1). Heatmaps, UpSet plots, and Volcano plots were generated with R v4.3.1 using the “pheatmap” (Kolde, 2019), “UpSetR” (Gehlenborg, 2019) and “ggplot2” (Wickham, 2016) packages, respectively.

### Metabolite analysis

Approximately 50 mg (fresh weight) of liquid nitrogen frozen and pulverized protonema tissue of WT, Δ*PpFLOT-1,* and all *PpFLOT*-OEX lines was mixed with 700 µL of basic acetone (acetone: 0.2 M NH_4_OH, 9:1 v/v) containing 2 µl corticosterone (2 mg/mL) per sample as an internal standard, in a 2 ml Eppendorf tube. Samples were stored at −20°C for 20 min with gentle mixing every 5 min, then centrifuged for 10 min at 4 °C at maximum speed. The supernatant was transferred to a new tube, and the pellet was extracted a second time using 300 µl of basic acetone. Both supernatants were combined, mixed, and 750 µL of the mixture was vacuum dried (Concentrator 5301; Eppendorf). Sample preparation was performed under low light, and the dried supernatant was stored at −80°C until further analysis, with argon added to prevent oxidation. The dry pellet was resuspended in 100 µl methanol, and subsequent LC-MS/MS was performed using a Dionex Ultimate 3000 UHPLC with a diode array detector (DAD) (Thermo Fisher Scientific). For pigment analysis, a 5 µl injection volume was separated at a flow rate of 500 µL min-1 on a C30 reversed-phase column (Acclaim C30, 3 µm, 2.1 x 150 mm, Thermo Fisher Scientific) at 15°C. ACN (solution A) and a mixture of methanol and ethyl acetate (50/50; v/v) (solution B) both containing 0.1% formic acid were used to form a solvents gradient. The gradient started with 14.5 % solution B followed by a ramp to 34.5 % solution B within 15 min that was then maintained for 10 min, before returning to 14.5 % solution B with additional 5 min of re-equilibration. Pigments were quantified by DAD. Analysis of the lipid composition was performed by separating 5 µl injection volume at a flow rate of 500 µL min^-1^ on a C30 reversed-phase column (Acclaim C30, 3 µm, 2.1 x 150 mm, Thermo Fisher Scientific) at 15°C. The solvent used for lipid separation was water (solution C) and an ACN:isopropanol mixture (7:3; v/v) (solution D), both including 1 % ammonium acetate and 0.1 % (v/v) acetic acid. The 26 min gradient started at 55% solution D, followed by a ramp to 99% solution D within 15 min. After a 5 min washing step at 99% solution D, the gradient was returned to 55% solution D and kept constant for 5 min equilibration. For lipid detection, an electrospray ionization (ESI) source was used in positive mode and negative mode to detect fatty acids. Nitrogen was used as the dry gas, at 8 L min^−1^, 8 bar, and 200°C. Mass spectra were recorded in MS mode from 50 m/z to 1300 m/z with 40.000 resolution, 1 Hz scan speed, and 0.3 ppm mass accuracy using the timsTOF (Bruker) mass spectra were recorded in MS mode. Compounds were compared to reference standards or annotated in a targeted approach using DAD data as well as the specific mass (m/z) at retention time and the isotopic pattern. Data were acquired by OTOF Control 4.0 and evaluated using DataAnalysis 5.0 and MetaboScape 4.0. All software tools were provided by Bruker. All analyses were performed in four replicates, and extreme outliers were removed while ensuring that at least three replicates per line remained for statistical analysis. To identify significantly enriched or depleted pigments and lipids, log_2_ transformation of the DAD data was performed, and the fold change compared to the WT was calculated. Subsequently, statistical analysis was conducted using Perseus v 2.0.1.1. (Tyanova et al., 2016; Tyanova and Cox, 2018). A two-sample Student’s *t*-test with was performed for all generated mutant lines using the WT as the control group. Metabolites were considered statistically significant when p < 0.05, and permutation-based FDR q < 0.05. All presented heatmaps and UpSet plots were generated using R v4.3.1 with the “pheatmap” (Kolde, 2019) and “UpSetR” (Gehlenborg, 2019) packages, respectively.

### TEM microscopy

Protonema cultures of WT, Δ*PpFLOT-1*, and all *PpFLOT*-OEX lines were harvested and fixed for three days in fixation buffer made of 75 mM cacodylate, 2mM MgCl_2_ (pH = 7.0) and 2.5 % glutaraldehyde. Subsequently, the samples underwent sequential washing steps in fresh fixation buffer for 5, 20, 40, and 60 min. Following this, samples were incubated for 90 min in fixation buffer supplemented with 1% OsO_4_, followed by washing in fresh buffer for 25 min. After overnight incubation in fresh buffer, the samples underwent additional washing steps in ddH_2_O for 5, 15 and 30 min. The samples were then gradually dehydrated by incubating them for 20 min in 10 % acetone, 20 min in a 20 % acetone/ 1% UrAc mixture, 110 min in 40 % acetone and for 15 min in 60 % acetone, 80 % acetone. Prior to embedding in 100 % resin, the samples were subjected to a final incubation in 100 % acetone in three steps. Ultrathin sections were prepared, and the slides were analyzed using a Zeiss EM912 (Carl Zeiss Microscopy GmbH) equipped with a 2k x 2k Tröndle slow-scan CCD camera (TRS, Tröndle Restlichtverstärkersysteme, Moorenweis, Germany) operated at 80 kV.

## Acknowledgments

This project was carried out in the framework of MAdLand (https://madland.science/, DFG Priority Program 2237). WF and EC are grateful for funding by the Deutsche Forschungsgemeinschaft (DFG; FR 1677/5-1). We also acknowledge help by Peter Geigenberger and David González-Gampo introducing the measurement of electrolyte leakage and technical help by Ioana Miruna Raulea.

## Author contributions

EC, OT and WF designed the research. EC and NN performed the research. SM and H-HK performed confocal microscopy and SS and ML performed LC-MS experiments. Transmission electron microscopy was performed by AK. EC, SS and ML performed bioinformatical analysis of proteomics and metabolomics data. Statistical analysis was performed by EC. All authors analyzed the data. EC, OT and WT wrote the manuscript. All authors reviewed the manuscript.

## Data availability statement

All data generated or analyzed during this study are included in this article and its supplementary information. The raw proteomics and metabolomics data will be open to the public at the proteomics identification database (PRIDE) after the acceptance of the manuscript.

## LIST OF SUPPLEMENTARY TABELS AND FIGURES

### SUPPLEMENTARY FIGURES

**Supplementary Figure 1:**
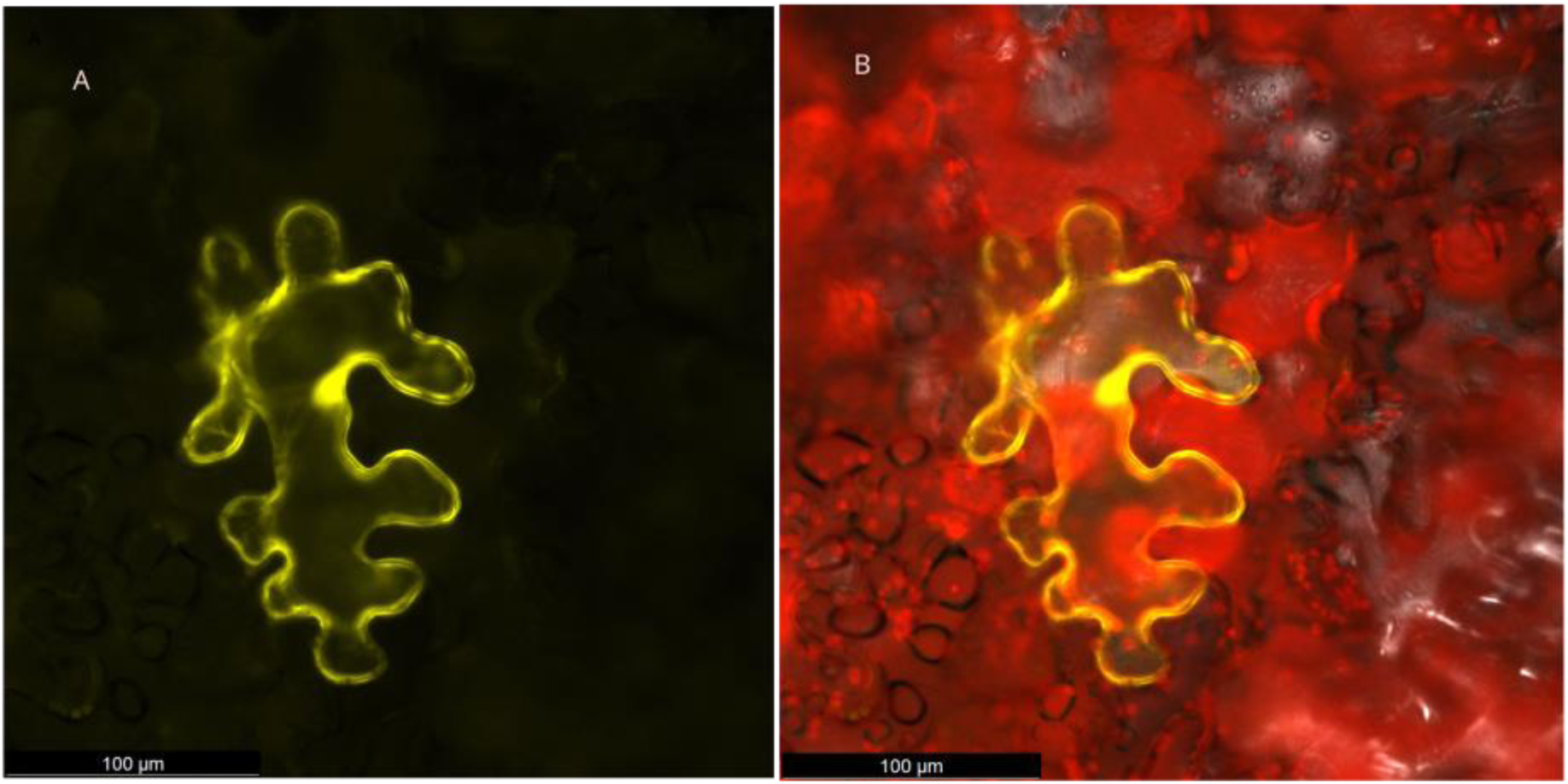
Confocal microscopy images of PpFLOT::venus in tobacco leaf cells showing PpFLOT::venus (A, yellow) and chlorophyll (B, red). *N. benthamiana* leaves. Scale bars are indicated in the respective images.

**Supplementary Figure 2:**
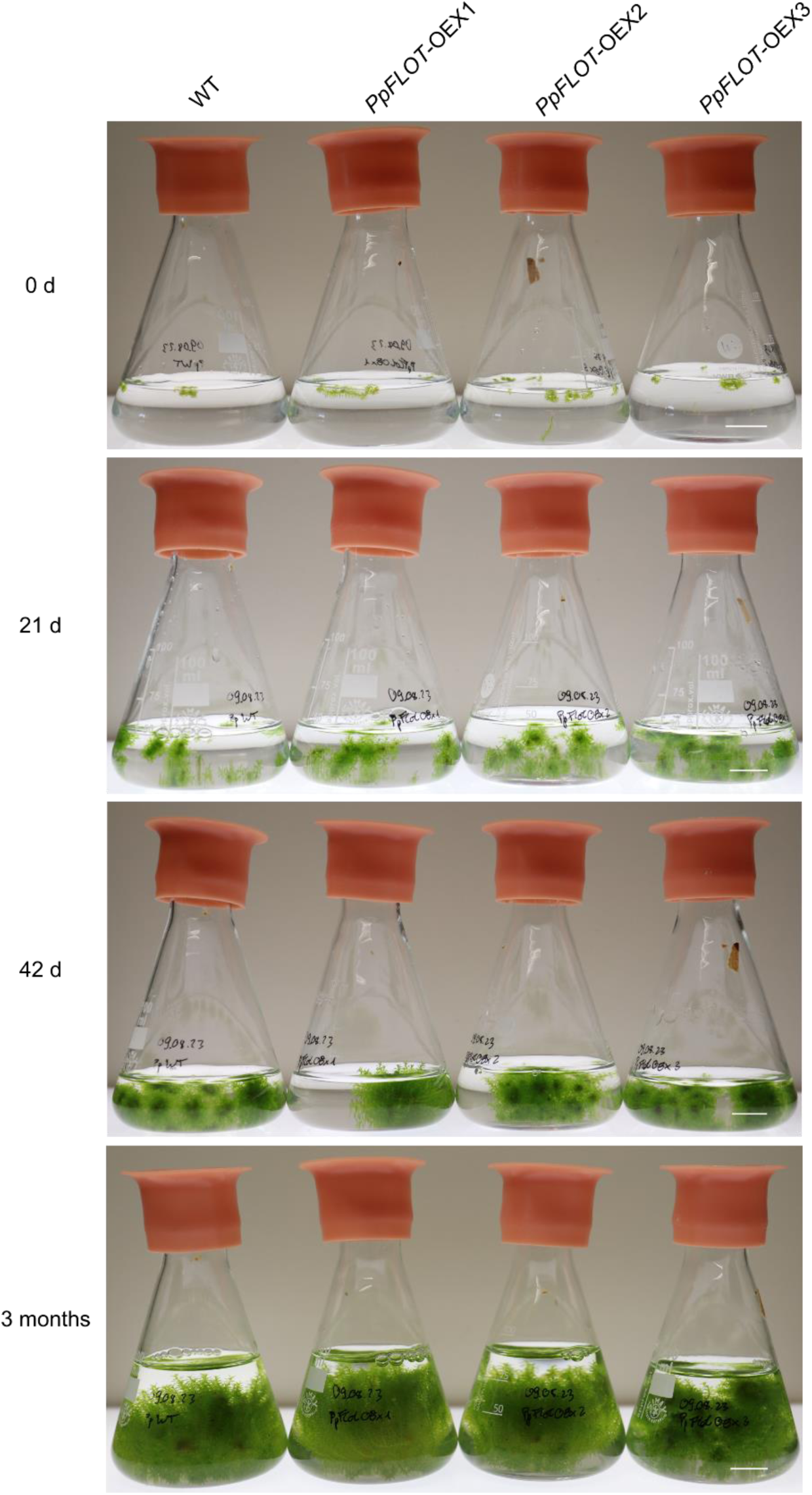
Growth of gametophores from *PpFLOT*-OEX lines submerged in liquid medium. *P. patens* WT and all three generated *PpFLOT*-OEX lines previously grown on solid medium were transferred to 50 ml liquid medium and observed for three months. Cultures were grown with continuous shaking at long day conditions (16 h light, 8 h dark; LD 16:8). Liquid media levels were maintained as needed to keep growing gametophore colonies submerged in the media. Images were captured at 0 d, 21 d, 42 d, and three months after transfer. Scale bar represents 1 cm.

**Supplementary Figure 3:**
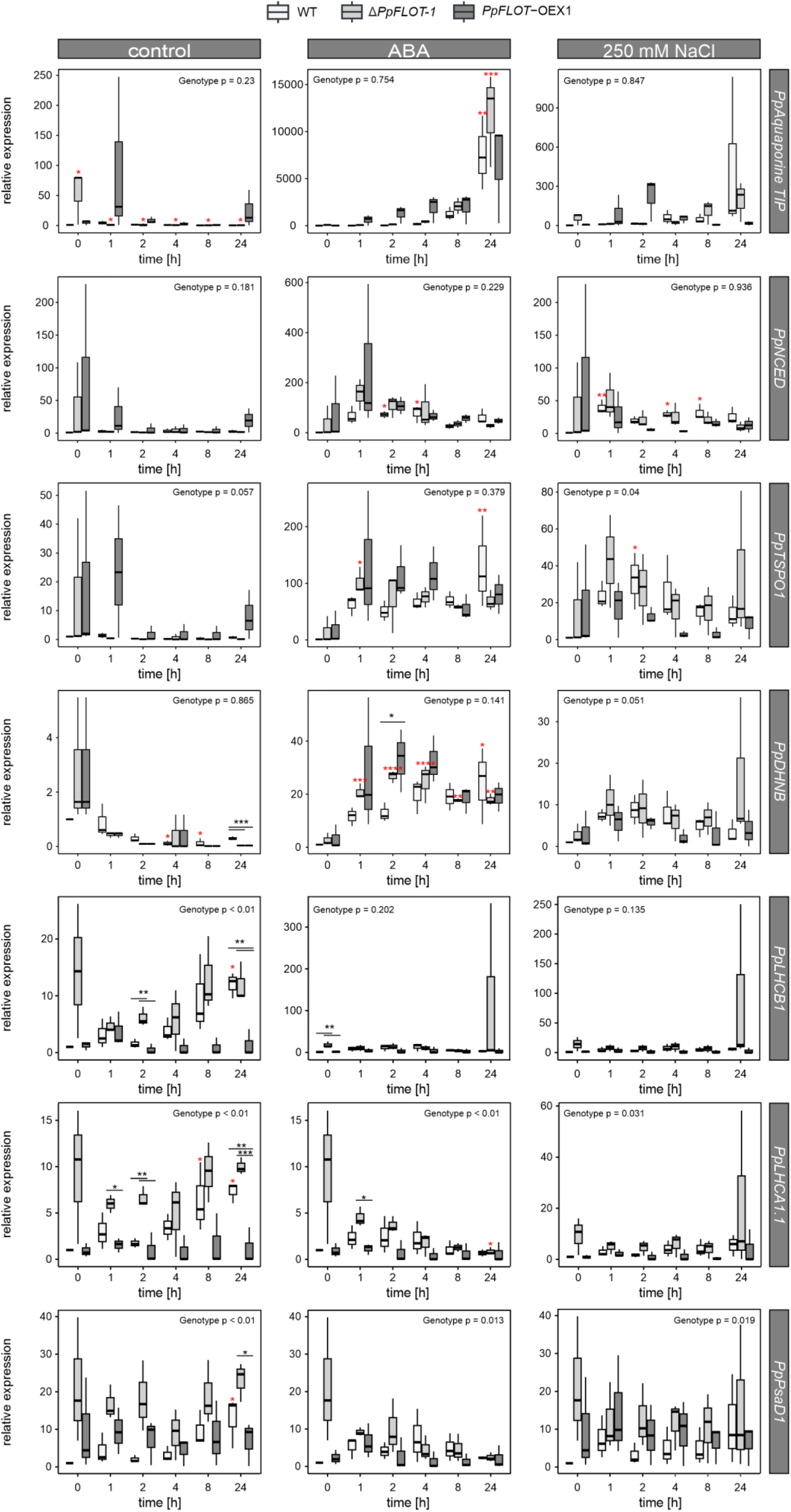
Relative gene expression analyses of salt-induced genes and genes encoding components of photosystem I and II under control conditions and after ABA and salt treatment. qRT-PCR results are shown as box blots illustrating the relative gene expression of *PpAquaporine TIP* (Pp1s44_31V6.1), *PpNCED9* (Pp1s412_7V6.1), *PpTSPO1* (Pp3c2_17540), *PpDHNB* (Pp3c5_11880), *PpLHCB1* (Pp3c2_35930), *PpLHCA1.1* (Pp3c13_14980) and *PpPsaD1* (Pp3c16_23780) compared to WT expression at 0 h and normalized against *PpEF1α* as described by Schmittgen and Livak (2008). Measurements were conducted over 24 h under control conditions (left) and treated with ABA (middle), and 250 mM NaCl treatment (right). The relative gene expression of the respective genes was measured for Δ*PpFLOT-1*, WT and *PpFLOT*-OEX1 protonema under control conditions in biological triplicates. Statistically significant changes in expression between the three genotypes was determined by ANOVA, with p-values provided in the respective graphs. Significant differential gene expression at a specific time point is indicated by black asterisks. Results of the Tukey’s HSD of the time-dependent expression within the same genotype are marked by red asterisks when significant compared to 0 h of treatment. * p < 0.05, ** p < 0.01, *** p < 0.001, **** p < 0.0001.

***Supplementary Table 1:*** Oligonucleotides used in this study.

***Supplementary Table 2:*** Significantly differentially accumulated pigments in all *PpFLOT* mutant lines compared to the WT.

***Supplementary Table 3:*** Results of ANOVA analysis comparing z-scores between WT and all *PpFLOT*-OEX lines.

***Supplementary Table 4:*** Results of the GO-term analysis of all differentially expressed protein groups between at least one tested genotype.

***Supplementary Table 5:*** Results of Student’s *t*-test comparing protein expression of all *PpFLOT*-OEX lines with the WT.

***Supplementary Table 6:*** Significantly differentially accumulated lipids in all *PpFLOT* mutant lines compared to the WT.

